# Predator adaptation to single prey species yields positive specialization and multiple forms of diversification

**DOI:** 10.64898/2026.05.15.725110

**Authors:** Marco La Fortezza, Sophie Scheiwiller, Sébastien Wielgoss, Yuen-Tsu Nicco Yu, Severin Frei, Gregory J. Velicer

**Affiliations:** Institute for Integrative Biology, Department of Environmental Systems Science, ETH Zurich, Zurich, Switzerland

## Abstract

Ecological specialization emerges when adaptation to a focal context increases fitness in that context relative to others. Experimental evolution has been widely used to study microbial specialization in abiotic environments but not predatory specialization. Here we demonstrate evolutionary specialization by a bacterial predator and characterize associated diversification of its predation profile. Populations of the bacterium *Myxococcus xanthus* evolving on single, non-evolving prey species diverged in predatory performance, showing increased performance on their home prey relative to their common ancestor, and relative to foreign prey not encountered during adaptation. Adaptation to the single-prey environments resulted in striking radiation of performance profiles across a diverse panel of foreign prey that was shaped interactively by selection, chance, and indirect effects, with home-prey identity modulating the degree of stochastic indirect diversification. Despite a great diversity of indirect evolutionary effects, correlated evolution was net-positive, yielding positive predatory specialization as the general outcome. Genomic evolution mirrored phenotypic evolution in that degrees of genomic parallelism differed as a function of home-prey identity. These findings show that adaptation to even simple biotic conditions can generate great ecological and behavioral diversity, linking direct selection, deterministic indirect effects of adaptation, stochasticity, and the origins of predator specialization and diversification.

## Introduction

Ecological specialization – the evolution of enhanced performance in particular ecological contexts relative to others – is a common outcome of adaptive evolution and a major source of biological diversity^1–3^. Specialization arises when adaptation to one selective context results in a magnitude of fitness increase in that context that is not matched by indirect fitness increases of equal magnitude in other contexts. Such specialization is thought to often result from fitness tradeoffs mediated by antagonistic pleiotropy^4,5^, but can also arise from hitchhiking of linked alleles and relaxed selection on unused traits^6–8^. Experimental evolution with microbes adapting to abiotic conditions has shown how such indirect evolutionary effects can diversify ecological profiles, exposing latent phenotypic variation that becomes manifested in new contexts^3,9,10^. However, how indirect evolutionary effects shape biotic interactions such as predation and associated specialization has been less explored experimentally^9,11–13^.

Predators exhibit striking variation in their abilities to kill and consume other organisms^14–17^. We define a predator’s ‘predation profile’ as its relative predatory performance across a specified set of potential prey organisms. Predators encounter a range of distinct organisms that might serve as their prey, each favoring a potentially different suite of behavioral, morphological and physiological predator traits. Adaptive improvement at predation on one prey type thus seems unlikely to fully carry over to other prey as a general rule, resulting in predatory specialization, and indeed might often reduce fitness on other prey. Yet despite the centrality of predation to the ecological character of biological communities^18,19^, the process of predatory specialization has to our knowledge yet to be directly characterized in any non-virus system^20^, and how predation profiles evolve and diversify is poorly understood. Previous evolution experiments with microbial predators and prey have examined adaptive evolution by prey and evolutionary change by the predator in the selective context of the experiment^21^. They have also investigated indirect effects of adaptation to defined prey environments on seemingly unrelated social traits^22^ and on interactions with bacteriophage^9^. However, they have yet to investigate how evolutionary adaptation to one prey type (or set of prey) fueled by spontaneous mutation affects performance across other prey types; indirect effects of predatory adaptation with respect to predator specialization and diversification remain unexplored.

Indirect evolutionary effects can generate three broad classes of specialization – negative, neutral or positive (Fig. 1a). Negative specialization occurs when adaptation in one ecological context (such as interaction with one prey type) reduces performance in other contexts, for example from antagonistic pleiotropy^23–25^, which has long been viewed as a major constraint on adaptive generalism^1–3,5,26,27^. Neutral specialization emerges when adaptation affects performance only in a focal selective context, with little effect on fitness in other contexts^8,13^. Positive specialization occurs when adaptation to one context improves average performance in others, but to a lesser degree than improvement in the adaptive context^9,13,26,28^. (Positive specialization can also be inversely viewed as incomplete generalist improvement (Fig. 1a).) Microbial evolution experiments in abiotic environments have revealed all three forms, often within the same system, even when there is an average trend toward either negative or positive specialization^13,24,26,28,29^. These categories comprise a continuum of evolutionary outcomes that connect mutation effects with an organism’s ecological performance profile.

**Figure 1.**
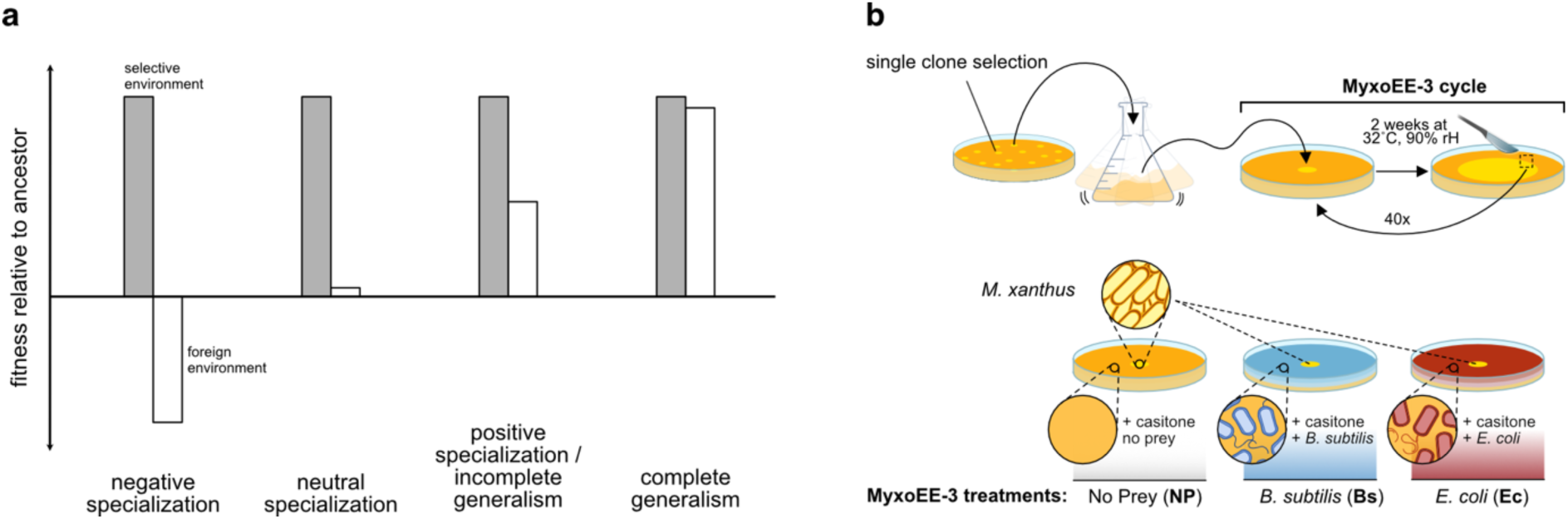
Types of evolutionary specialization and MyxoEE-3 procedure and treatments. **a)** Categories of specialization/generalism differing according to the character of indirect effects of evolution in a particular selective context. Grey bars indicate increases in fitness by a lineage/population in the focal selective environment compared to its ancestor, while white bars show indirect fitness change in other foreign environments not encountered during selection. Indirect effects can be considered with respect to a single foreign environment or averaged across multiple foreign environments. Positive specialisation can inversely be understood as incomplete generalism. **b)** MyxoEE-3 cycling and treatments. The top image shows the initiation and cycling scheme common to all MyxoEE-3 treatments (adapted from La Fortezza *et al.* 2022). Replicate populations within MyxoEE-3 treatments were started from subclones of the common *M. xanthus* ancestor GJV1 (here collectively ‘Anc’, see Methods: *MyxoEE-3;* Rendueles *et al.* 2015 and La Fortezza *et al.* 2022). *M. xanthus* populations (yellow circles) allowed to swarm to their ability for two weeks before a rectangle from the most distal position on the swarm perimeter was cut and transferred to the center of a new plate to initiate the next cycle. Populations examined in this study evolved over 40 two-week cycles. The bottom image shows the distinguishing features of the three MyxoEE-3 treatments examined here, namely CTT agar (1% casitone, 1.5% agar) without prey (No Prey, **NP**), CTT agar overgrown with a lawn of *B. subtilis* (blue, **Bs**), or CTT agar overgrown with a lawn of *E. coli* (red, **Ec**).

Myxobacteria (order Myxococcota), including the model species *Myxococcus xanthus,* are powerful systems for directly studying predator evolution, including specialization and diversification of predator profiles, with experimental evolution^21^. They inhabit a broad range of soil and aquatic environments and can kill and consume a wide range of microbial prey, including gram-negative and gram-positive bacteria and various eukaryotic microbes^30–33^. *M. xanthus* is a social predator that “searches” for prey^21,34,35^ using collective motility^36–39^, and “handles” its prey by delivering lethal and lytic compounds both via direct contact with prey cells^39,40^ and remotely with secreted compounds^40,41^. Beyond predation, myxobacteria exhibit highly social life-cycles that include aggregative development of spore-bearing multicellular fruiting bodies under starvation^42^ and cooperative spore germination^43^.

Natural isolates of *M. xanthus* differ greatly in their predation profiles, for example in their abilities to swarm through single-species lawns of diverse prey species^32,33,44^ while killing and consuming them in the process^44^. This variation suggests that prey-driven selection may be an important source of predatory diversity in nature. We investigated how such diversity might evolve from a single evolutionary starting point under controlled conditions both due to direct responses to selection imposed by multiple highly diverged prey species and from indirect effects of evolution while consuming those prey. Specifically, we used *M. xanthus* populations from the evolution experiment MyxoEE-3^11,22,45,46^, in which replicate predator populations were propagated for hundreds of generations on agar surfaces containing non-evolving lawns of either *Bacillus subtilis* or *Escherichia coli,* or on nutrient agar without prey, while the predators were under selection for increased fitness at the front of expanding swarms (Fig. 1b). We then measured how adaptation to those home-prey environments affected proxies of predatory fitness, both on their home prey and across a diverse panel of foreign prey not encountered during MyxoEE-3. This design allowed us to characterize predator diversification due to the individual and interactive effects of adaptation to different home prey, indirect evolutionary effects on performance with foreign prey, and chance.

## Results

### Adaptation corresponds with increased swarming in home environments with prey

We used two approaches to investigate adaptation by evolved populations to their respective MyxoEE-3 home selective environments after 40 cycles. First, we compared the swarming rates of evolved populations in their home environment to the swarming rates of their ancestral subclones (Supp. Fig. 1a), hereafter collectively referred to as ‘the ancestor’. During *M. xanthus* growth on prey lawns, swarming rate has been found to correlate with both total prey killed and predator growth, showing this trait to be an important component of overall predatory performance in these environments^32,44,47,48^; it was thus expected to be a likely correlate of adaptation in MyxoEE-3^49^ (see Methods: *MyxoEE-3* and *Swarming assay*).

The ancestor swarmed at different rates on the three MyxoEE-3 substrates featured in this study, being fastest in the absence of prey (CTT nutrient agar only), and faster through *E. coli* lawns than through *B. subtilis* lawns(Fig. 2a). The latter outcome is consistent with previous studies indicating that *E. coli* fuels more growth by *M. xanthus* than do *Bacillus* species^32,44,48^.

**Figure 2.**
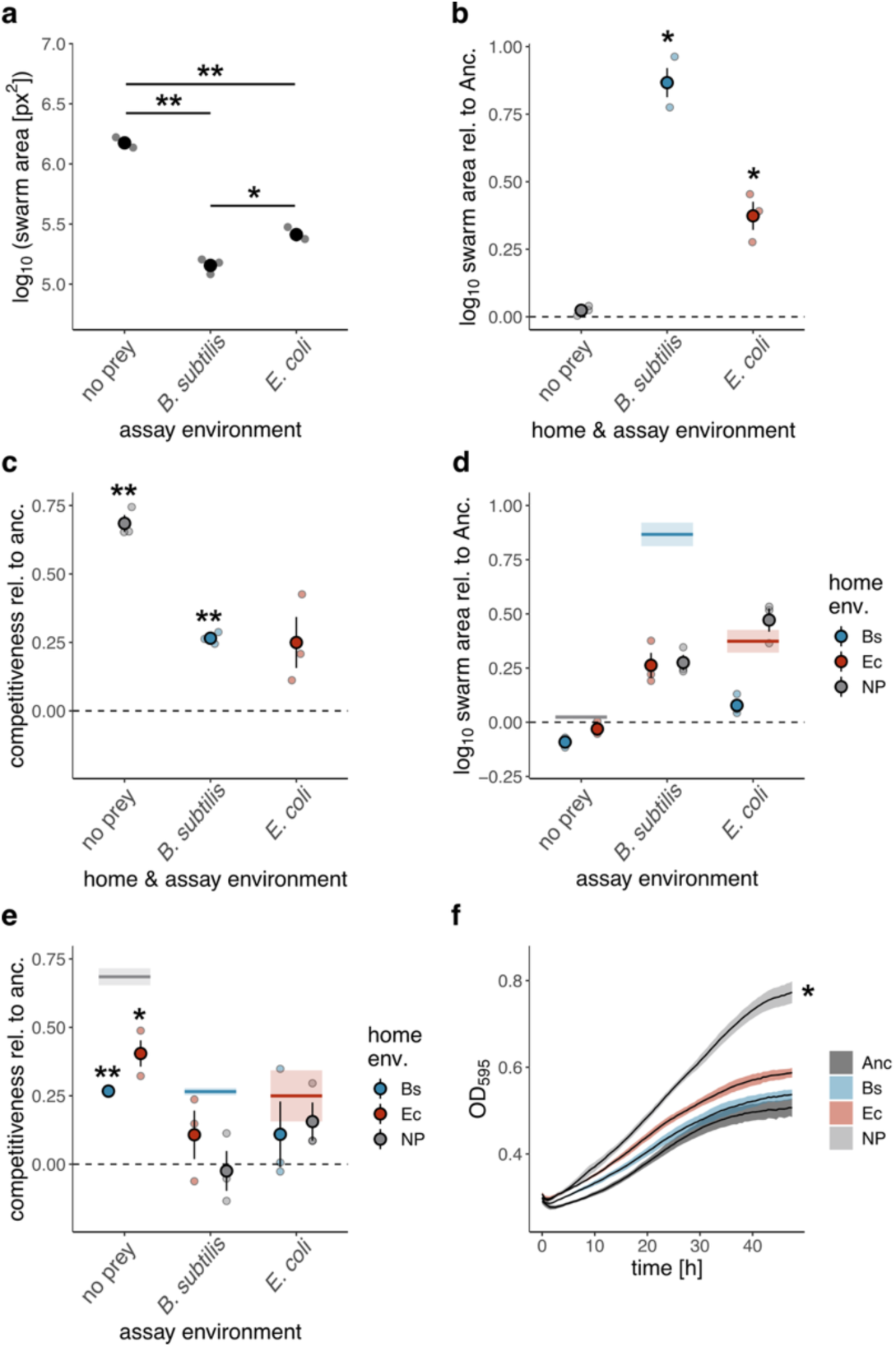
Adaptation to MyxoEE-3 home prey resulted in specialized predatory performance without tradeoffs toward the foreign MxyoEE-3 prey species. **a)** Average swarm areas of the ancestral clones in the three MyxoEE-3 environments examined in this study. **b)** Average swarm areas of evolved populations in their home environment – the environment in which they evolved during MyxoEE-3 – relative to the ancestor (indicated by the dashed line at zero). Transparent dots indicate averages of evolved populations for the specified evolutionary treatment within a given assay replicate; opaque dots give cross-replicate averages (*n =* 3) (also for panels c, d and e). *x*-axis categories indicate the MyxoEE-3 environment in which the reported assay was conducted (also for panels c, d and e). **c)** Competition assays in home MyxoEE-3 environments: Evolutionary improvement by evolved populations at limiting the end-of-competition population size of the marked ancestor in 1:1 mixed culture in the home environment of the evolved populations. Values at the dotted line would indicate zero evolutionary change in this fitness proxy (also for panel e). **d)** Average swarm areas relative to the ancestor for each treatment in foreign MyxoEE-3 environments. The horizontal lines and associated shaded areas report the mean and SEM in the home environment (same values as in panel b). **e)** Competition assays in foreign MyxoEE-3 environments: Evolutionary improvement by evolved populations (or lack thereof) at limiting the end-of-competition population size of the marked ancestor in 1:1 mixed culture in the MyxoEE-3 environments examined in this study in which the specified evolved populations did not evolve. To facilitate comparisons, the horizontal lines and associated shaded coloured boxes report the mean values and SEM plotted in panel c. **f**) Growth curves measured in CTT liquid media, averaged across replicate evolved populations for each MyxoEE-3 treatment. The shaded areas show the associated SEM for three assay replicates. Error bars indicate SEM across biological replicates. * indicates *p* < 0.05, and ** indicates *p* < 0.01 for significant differences between environments or between the ancestor and evolved populations where appliquable (also for other panels); see Data Source for all calculated *p* values.

Consistent with previous results^49^, the four populations that evolved on CTT nutrient agar without prey (NP for ‘no prey’) showed only a small and insignificant increase in swarming rate in their home environment (*p* = 0.160, one-sample *t*-test, n = 3, df = 2; Fig. 2b, Supp. Fig. 1d). In contrast, the populations evolved on *B. subtilis* (Bs) or *E. coli* (Ec) increased greatly in swarming rate in their respective home environments, with the Bs-evolved populations increasing the most (swarm areas 7.5-and 2.4-fold greater than the ancestor, respectively, *p*_Bs_= 0.020; *p*_Ec_= 0.038, one-sample *t*-tests, n = 3, df = 2; Fig. 2b, Supp. Fig. 1d). Thus, the average degree of evolutionary increase in swarm area across the three MyxoEE-3 environments examined here correlated inversely rank-wise with the swarming rate of the ancestor in those environments (Fig. 2a, b). All four Bs populations increased in their home-environment swarming rate, while three of the four Ec populations did so (Supp. Fig. 1d).

In our second approach, we tested for adaptation more directly by performing mixed-culture competition experiments. In these competitions, we compared growth of a genetically marked variant of the ancestor when it was mixed 1:1 with evolved populations in their home environment relative to when it was mixed with the unmarked ancestor, its parent (see Methods: *Competition assay*, Supp. Fig. 1b). As expected, the evolved populations from each of the three MyxoEE-3 treatments showed increased competitiveness, on average, against the marked ancestor in their home environment, clearly for NP and Bs and suggestively for Ec (Fig. 2c, Supp. Fig. 1e; NP: 68% increase, *p*_NP_ = 0.002; Bs: 27% increase, *p*_Bs_ = 0.002; Ec: 25% increase, *p*_Ec_ = 0.115; one-sample *t*-tests, n = 3, df = 2). In other words, in their home environments, evolved populations tended to reduce growth by the marked ancestor compared to its growth in mixture with the unmarked ancestor. Increased competitiveness was greater in magnitude for NP populations than Bs and Ec populations, which increased to similar degrees.

Comparing the home-environment competition results (Fig. 2c) to the swarming results (Fig. 2b) indicates that increased home competitiveness by the NP lines was not associated with increased home swarming, a result consistent with a prior study^49^ and suggestive of adaptation by interference competition. In contrast, populations adapted to prey increased in both swarming rate and competitiveness, suggesting different mechanisms of adaptation at play in the presence vs absence of prey.

### Home-prey adaptation results in positive specialization

We also used competition experiments to ask whether evolved increases in competitiveness in home environments would fully transfer to foreign MyxoEE-3 environments (complete generalist improvement) or would be associated with negative, neutral or positive specialization (Fig. 1a). The three treatments of evolved populations differed in their patterns of coincidental evolution of competitiveness. The NP populations showed no and little carryover of their large home-environment increases in competitiveness to the *B. subtilis* and *E. coli* environments, respectively (Fig. 2d). In contrast, the Bs and Ec evolved populations showed complete generalist improvement, *i.e.* increased competitiveness in the no-prey environment of a magnitude similar to or greater than their respective home-environment increases. This generalist indirect effect of adaptation to prey on no-prey competitiveness cannot be attributed to adaptation to shared culture conditions because the same generalist effect was not observed in the opposite direction. The Bs and Ec evolved populations increased in competitiveness on the alternative MyxoEE-3 prey only insignificantly (Fig. 2d), and to a lesser degree than their home-competitiveness increased, suggesting either weakly positive or neutral specialization.

Because the NP populations had only casitone as a carbon source to fuel growth during MyxoEE-3 while the *Bs* and *Ec* populations could consume *B. subtilis* or *E. coli,* respectively, as well as potentially any residual casitone not consumed by the prey, we expected more adaptation to growth on CTT alone by the NP populations than by the Bs or Ec evolved populations. To test this, we measured population growth in liquid CTT over 48 hours. Growth rate in CTT liquid increased in all evolutionary treatments, but did so only little among the Bs populations, slightly more in the Ec populations, and most, as expected, in the NP populations (Fig. 2e, *p*_NP_ = 0.018, paired *t*-test, n = 3, df = 2). Bs populations varied the most in their growth dynamics, with Bs-4 growing the fastest and Bs-1 having actually decreased in growth rate, despite showing increased swarming (Supp. Fig. 1d, 1f). Thus, increased efficiency at using casitone as a resource appears to explain a greater proportion of the adaptation shown by the NP lines – which were fed only casitone – than of the adaptation by the Bs or Ec lines.

Our analysis of swarming in home environments and competitiveness in both home and foreign MyxoEE-3 environments show substantial diversification of predatory profiles across the twelve examined populations relative to the ancestral clones due to both chance and selection. Within each of the NP, Bs and Ec treatments, replicate populations diversified by chance in both their swarming rates and competitiveness in their home selective environment (Supp. Fig. 1d, e). Populations also diversified at the treatment level in their competitiveness profiles across the three MyxoEE-3 environments due to selection.

### Home-prey adaptation greatly diversifies predator profiles across a broad panel of foreign prey

We also examined how adaptation to MyxoEE-3 environments might have resulted in specialisation and diversified *M. xanthus* predatory swarming profiles more broadly. We chose a phylogenetically diverse panel of 12 prey not used in MyxoEE-3, including six gram-positive and six gram-negative species. The selected foreign prey varied greatly in their apparent susceptibility to predation by the *M. xanthus* ancestor, including species through lawns of which the predator could efficiently swarm (*e.g. Allorhizobium vitis* and *Curtobacterium citreum*) and others on which predatory swarming was very poor or absent (*e.g. Priestia megaterium* and *Pseudomonas tolaasii*) (Table 1).

**Table 1:**
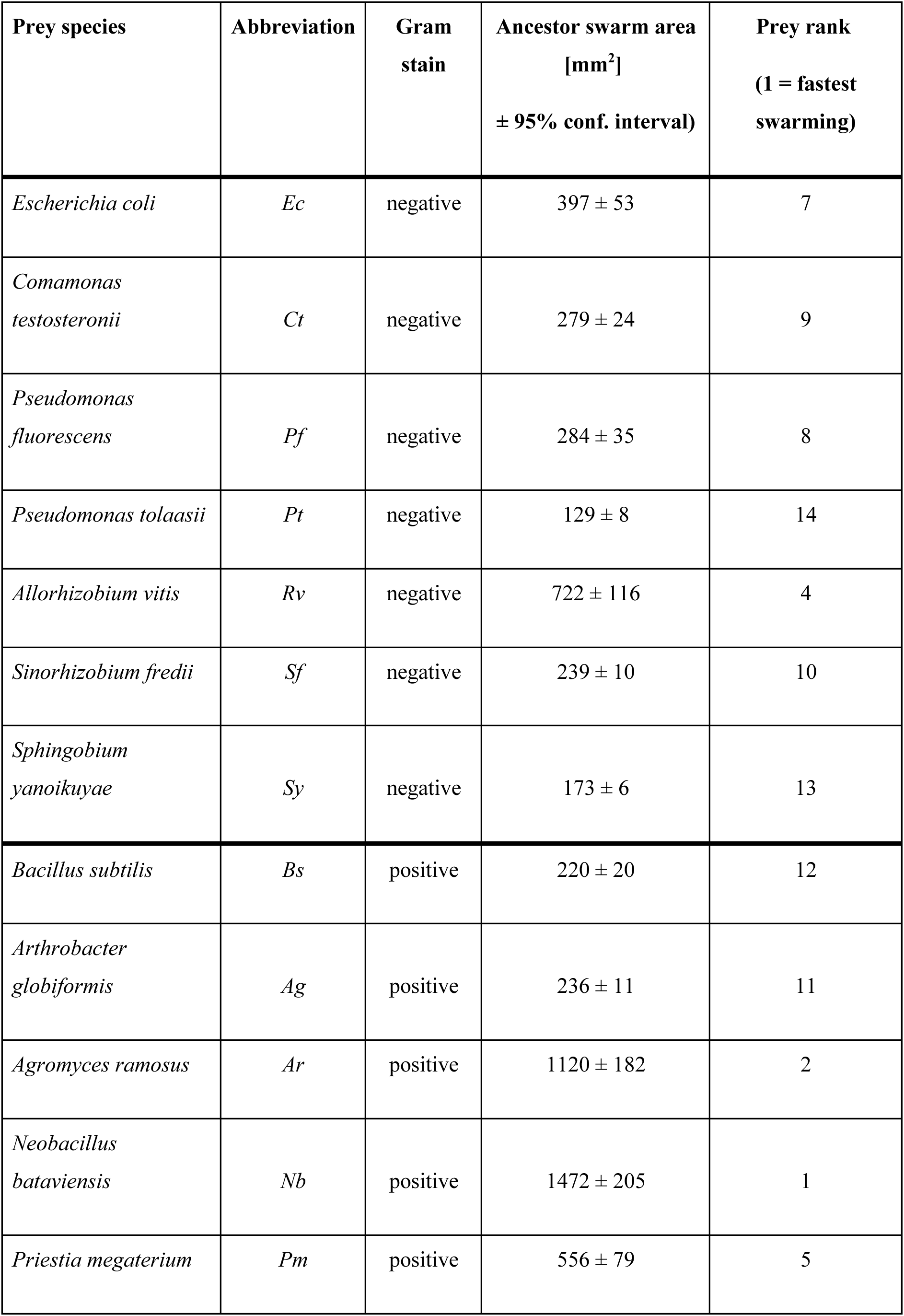
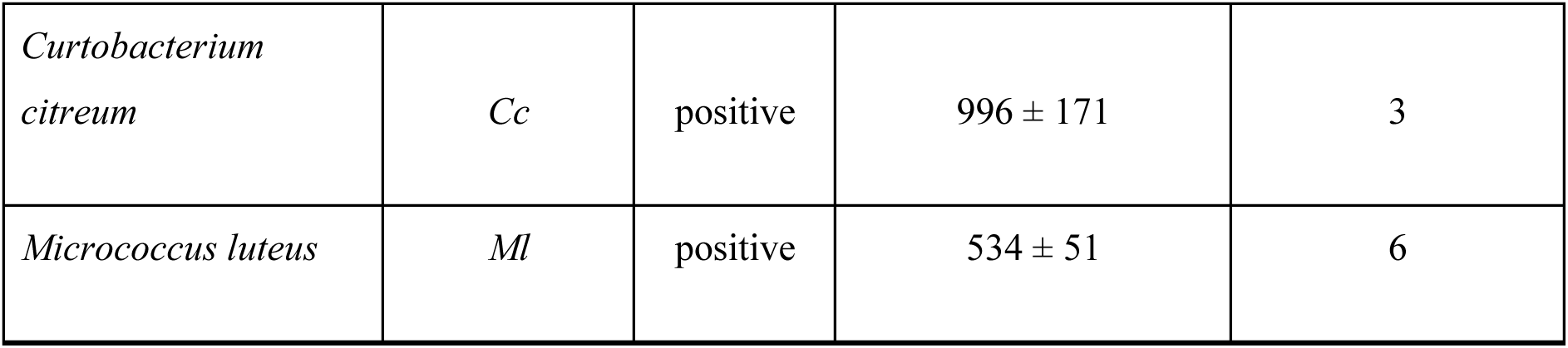
Prey species, their Gram stain and prey quality for the ancestor.

| **Prey species** | **Abbreviation** | **Gram stain** | **Ancestor swarm area [mm<sup>2</sup>]**<br>**± 95% conf. interval)** | **Prey rank**<br>**(1 = fastest swarming)** |
| --- | --- | --- | --- | --- |
| *Escherichia coli* | *Ec* | negative | 397 ± 53 | 7 |
| *Comamonas testosteronii* | *Ct* | negative | 279 ± 24 | 9 |
| *Pseudomonas fluorescens* | *Pf* | negative | 284 ± 35 | 8 |
| *Pseudomonas tolaasii* | *Pt* | negative | 129 ± 8 | 14 |
| *Allorhizobium vitis* | *Rv* | negative | 722 ± 116 | 4 |
| *Sinorhizobium fredii* | *Sf* | negative | 239 ± 10 | 10 |
| *Sphingobium yanoikuyae* | *Sy* | negative | 173 ± 6 | 13 |
| *Bacillus subtilis* | *Bs* | positive | 220 ± 20 | 12 |
| *Arthrobacter globiformis* | *Ag* | positive | 236 ± 11 | 11 |
| *Agromyces ramosus* | *Ar* | positive | 1120 ± 182 | 2 |
| *Neobacillus bataviensis* | *Nb* | positive | 1472 ± 205 | 1 |
| *Priestia megaterium* | *Pm* | positive | 556 ± 79 | 5 |
| *Curtobacterium*<br>*citreum* | *Cc* | positive | 996 ± 171 | 3 |
| *Micrococcus luteus* | *Ml* | positive | 534 ± 51 | 6 |

Adaptation to the two MyxoEE-3 prey environments (Fig. 2) resulted in positive specialization overall, *i.e.* faster swarming on average across the foreign prey species (Fig. 3a) but with smaller increases than on home prey. However, the selective prey environments differed in their indirect effects in multiple respects. First, the average coincidental increase of the Bs populations across the 13 foreign prey was higher than the corresponding average increase for the Ec populations (and higher than the NP populations as well; *p*_Ec-Bs_= 0.014, *p*_NP-Bs_= 0.017, *t*-tests with Benjamini-Hochberg multiple-testing correction).

**Figure 3.**
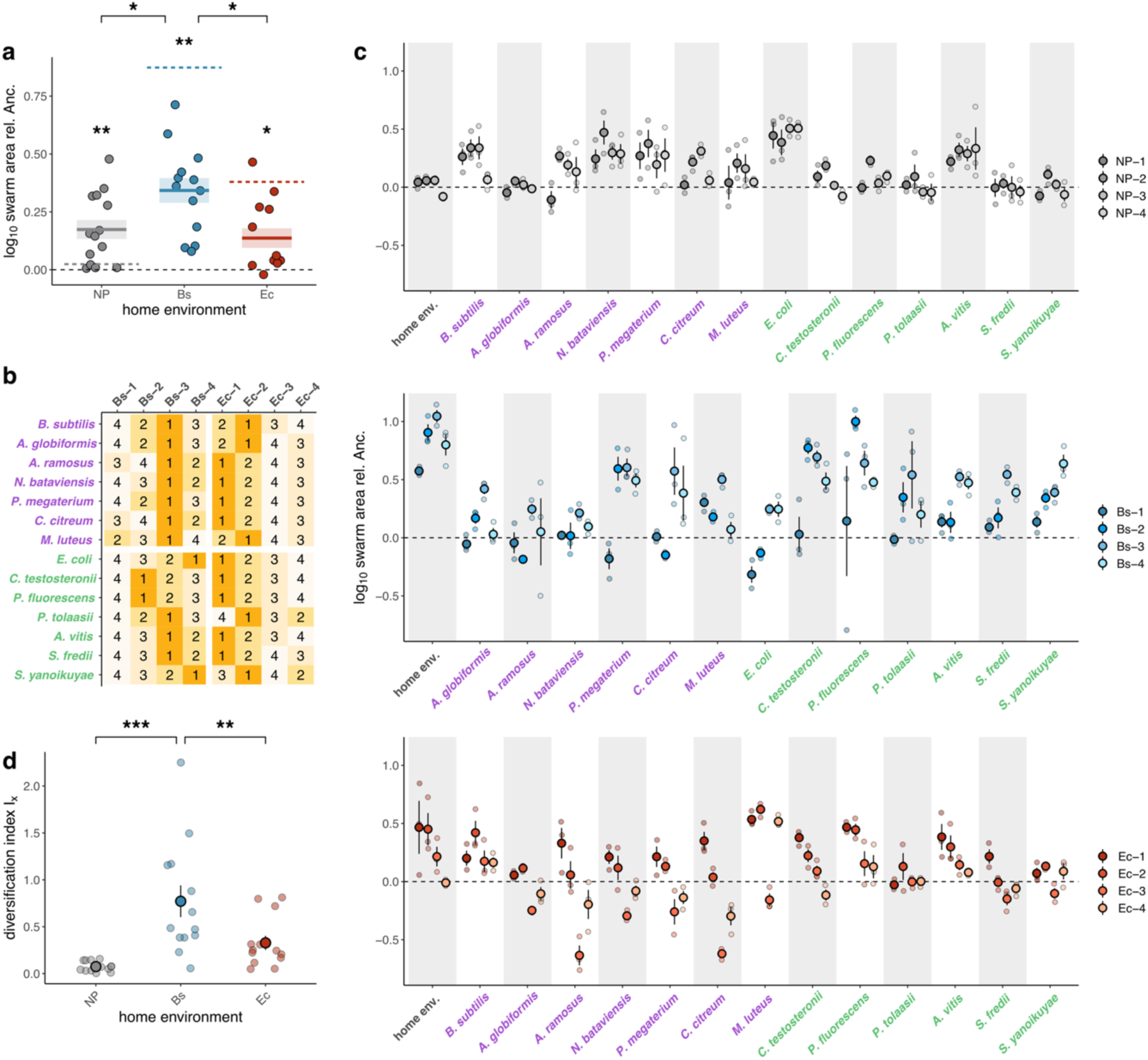
Adaptation to single-prey environments resulted in positive specialization and greatly diversified predatory-swarming profiles across a phylogenetically diverse panel of prey. **a**) Average proportional change in swarming rate of evolved populations categorized by MyxoEE-3 selective environment (NP, Bs and Ec) on 14 (NP populations) or 13 (Bs and Ec populations) foreign prey species relative to the ancestor. Zero (dashed black line) indicates no evolutionary change (also for panel c). Each dot represents the treatment-level average proportional change in swarm size for one foreign prey species. Colored dashed lines indicate the average swarming rate of the four populations for a given treatment in their home environment. Colored solid lines indicate average treatment-level swarming rates across all foreign prey examined; associated error bars (SEM, shaded boxes) reflect the degree to which proportional changes in evolved swarming (relative to the ancestor) vary across prey species. Asterisks indicate significant differences from 0, *i.e.* significant evolutionary change (* *p* < 0.05, ** *p <* 0.01). **b)** Ranks of coincidentally evolved swarming rates among evolved Bs and Ec populations for each foreign prey species examined; ‘1’ and ‘4’ indicate the fastest and slowest populations on a given prey type, respectively. **c**) Swarm sizes of individual evolved populations relative to the ancestor on lawns of foreign prey species; Gram-positive and -negative species names are given in purple and green, respectively. Large dots indicate the average values of three independent biological assay replicates (*n* = 3, reported as small dots) with the associated SEM. All *p* values are reported in Data Fig. 3. **d)** Diversification index *I_x_* calculated from the variance of the four replicate populations in each treatment swarming on 13 (Bs and Ec populations) or 14 (NP populations) foreign prey types adjusted by the average relative swarm area on each prey (following the calculation in Rendueles et al. 2017). The large dots with error bars represent the treatment average with the associated SEM. ** *p* < 0.01, *** *p* < 0.001

Second, treatments differed in the degree to which their replicate populations diversified in predatory swarming profiles by chance. Utilizing a previously developed diversification index (*I_x_*)^49–51^, we quantified within-treatment diversification in two ways. We first calculated diversification among replicate populations per prey type, averaging those 13 per-prey values (Fig. 3d). We also calculated diversification across the 13 prey per evolved population, averaging those per-evolved-population values (Supp. Fig. 3). With both approaches, the diversification index calculated for the Bs populations was higher than for the Ec and NP populations, with the differences in the first approach being highly significant (*p*_Ec-Bs_= 0.008, *p*_NP-Bs_< 0.001, *t*-tests with Benjamini-Hochberg multiple-testing correction). Thus, selection can modulate the role of chance in shaping the indirect diversification of organisms, an outcome also seen in a previous MyxoEE-3 study^51^.

As a specific example of indirect within-treatment diversification, populations Bs-1, Bs-2 and Bs-4 showed no or negative change on one or more foreign prey while Bs-3 increased more systematically on all foreign prey (Fig. 3c, Supp. Fig. 2c). Among the Ec populations, two increased in swarming on their home prey either little or not at all (Ec-3 and Ec-4, respectively) and these tended to swarm the least across the foreign prey panel (Fig. 3c). The prey type also shapes how indirect evolution phenotypes manifest. Two populations which show little increase on their home and foreign prey otherwise, increased swarming on *M. luteus* especially (Bs-1 and Ec-4, Fig. 3c).The ranks of swarm expansion on foreign prey are consistently high or low for some populations (Bs-3 and Ec-3, resp.), but also frequently reverse among replicate populations as a function of prey type, highlighting the role of chance in diversification of predation profiles (Fig. 3b). Intriguingly, although the NP populations evolved without prey did not increase swarming in their home environment (Fig 2b, Supp. Fig. 1d), they nonetheless increased swarming on foreign prey (Supp. Fig. 2c), showing that adaptation to abiotic ecological factors can have large and unexpected effects on predation profiles.

Altogether, we see that evolution of twelve initially identical populations of *M. xanthus* in three MyxoEE-3 selective regimes profoundly diversified their predatory performance at both treatment and population levels, *i.e.* due to both selection and chance. They did so in several respects – swarming and competitiveness in their home environments, competitiveness in home vs foreign MyxoEE-3 environments, and swarming through many non-MyxoEE-3 foreign prey, including selection-determined diversification in the degree of indirect swarming-profile diversification caused by chance.

### Predator genome evolution

A previous study with *M. xanthus* and *E. coli* showed that predator-prey interactions can impose selection on both predators and prey that strongly influences genome evolution^52^. Here we investigated whether the presence and identity of non-evolving prey in MyxoEE-3 impacted predator-genome evolution by sequencing the metagenome of each evolved population and comparing it both with the ancestral genome and with other populations.

#### General evidence of selection

Across all populations, non-synonymous single nucleotide variant (SNV) mutations outnumbered synonymous SNVs and exceeded all other mutational types (240 mutations total; 82 polymorphic and 158 fixed; Supp. Fig. 4a, b; Supp. Table 2). Consistent with strong and widespread positive selection during experimental evolution, genome-wide dN/dS ratios calculated from the 124 fixed coding SNVs (Supp. Table 2) were ≥1.5 in all populations^53,54^. For all subsequent analyses, we considered all 240 mutations present at frequencies higher than 0.05 in their respective populations (see *Methods: Genomic data analysis)*.

Non-random parallel evolution at a given genetic locus (or set of functionally related loci) among independently evolving populations reflects the operation of selection^55^. In MyxoEE-3, parallel evolution might be distributed across multiple treatments, reflecting shared selective factors, or might be biased toward a subset of treatments, indicating selective features unique to that subset. For the twelve MyxoEE-3 populations considered here (four in each of three treatments), any case of two or more populations having mutated independently at the same gene or operon is unexpected by chance (*p =* 0.029 and *p* = 0.050 for genes and operons, respectively; see *Methods: Mutational probabilities*). Considering our set of 240 mutations, 32 genes and 35 operons were mutated in two or more of the twelve MyxoEE-3 populations examined here (∼20% and ∼25% of all mutated genes and operons, respectively (Supp. Fig. 4e). Seven percent (7%) of all individual mutations were present in more than one population (Supp. Table 4, Supp. Fig. 4e). The probabilities of a gene or operon being mutated in three or more populations are *p =* 0.0004 and *p =* 0.0011, respectively. Operons and genes mutated in three or more of the twelve evolved populations are shown in Figs. 4c and 4f, respectively.

**Figure 4.**
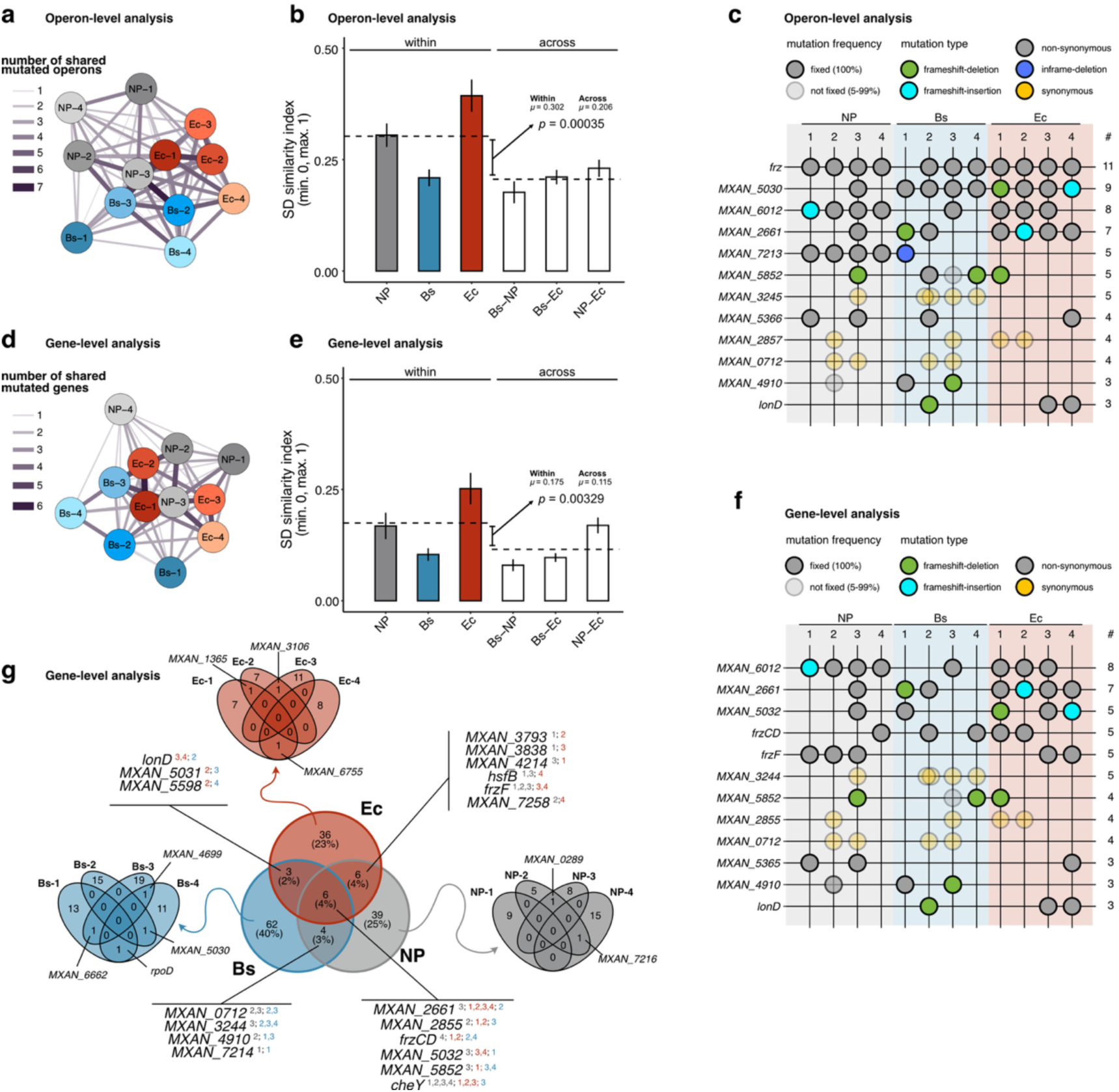
Prey-specific parallel evolution at the operon and gene level. *Operon-level analysis:* **a)** Network based on the number of mutated operons shared among evolved populations. Connector thickness and transparency are scaled to the number of shared mutated operons. Grey, blue and red nodes represent populations evolved in the NP, Bs and Ec treatments, respectively. **b**) Operon-level Sørensen-Dice (SD) similarity coefficients for pairwise comparison of populations within treatments and for each pairwise treatment combination. Average SD values for within- and between-treatment population pairs (coloured and white bars, respectively, *n* = 6 and 16 for within- and between-treatment averages, respectively; error bars indicate SEM) were calculated for comparison of similarity within and across treatments (dashed lines). The p-value indicates a significant difference between the overall SD averages for within- vs between-treatment comparisons (also for panel **e**). **c**) Operons with mutations in at least three of the 12 MyxoEE-3 evolved populations examined in this study. Dots indicate mutation presence for the respective population, and dot colour indicates the mutation type (see within-panel legend). Operon names were assigned based on the first gene present in each specific operon on the sense or antisense strand. *Gene-level analysis:* Panels **d), e)** and **f)** show the same analyses at the gene level as shown in panels a), b) and c) at the operon level. **g**) Venn diagrams reporting the number of mutated genes unique to vs shared between individual treatments (tri-colored diagram, centre), and unique to vs shared between individual replicate populations within treatments (monochromatic diagrams, margins). Genes mutated in more than one treatment (center) or more than one population within a treatment (margins) are identified by connector lines and gene names or *MXAN* numbers. For inter-treatment comparisons, superscripted small numbers identify the treatment and population-replicate in which the mutations occurred.

The *frz* operon was targeted most broadly and strongly by selection; it was mutated in all populations except Bs-1 (Fig. 4c), indicating that it facilitated adaptation to all three MyxoEE-3 environments. The *frz* operon is composed of six genes that collectively encode a protein complex involved in chemotactic responses and cell reversal during motility^56^. Of the six *frz* genes, only three were mutated: *frzCD, frzF,* and *frzE* (Supp. Table 4).

#### Treatment-specific genome evolution

We used several approaches to investigate potential differences in the character of parallel evolution at the treatment level: i) generation of “interaction networks”, ii) comparison of Sørensen-Dice (SD) similarity coefficients within vs between treatments, iii) operon-specific tests for non-random mutation distributions across treatments, and iv) treatment-specific tests for non-random mutation distributions across gene-ontology (GO) categories. We began by generating interaction networks in which a connecting line between two populations indicates at least one case of both populations having mutated the same gene or operon, with the length and thickness of connecting lines proportional to the number of such cases (see *Methods: Data analysis*). In such networks, spatial clustering of populations from the same treatment reflects treatment-specific evolutionary similarities, and indeed such clustering is apparent at both the operon and gene levels (Fig. 4a, d).

Next, we used the Sørensen-Dice (SD) similarity coefficient to quantify average degrees of genome-evolution similarity between i) replicate populations within each treatment and ii) paired treatments (Fig. 4b, e; see *Methods: Data analysis*)^57^. Parallel evolution as reflected by the SD coefficient was significantly higher within treatments than across paired treatments at both the operon and gene levels (randomisation test adapted from Deatherage *et al.* 2017, see *Methods: Data analysis*; *p*_operon_ = 0.00035; *p*_genes_ = 0.00329; Fig. 4b, e), indicating treatment-level genomic differentiation.

Within-treatment SD coefficients differed between treatments. Most notably, the four Ec populations evolved more similarly to one another at both gene and operon levels than did the Bs populations (Fig. 4b, e), which had within-treatment SD coefficients comparable to inter-treatment SD values. Considered inversely, the Bs populations diversified by chance more than did the Ec populations. The large difference in the SD coefficients of the two prey environments suggests that *M. xanthus* populations offered *B. subtilis* as prey had a more diverse range of adaptive genetic pathways readily accessible during MyxoEE-3 than the populations offered *E. coli*.

For three operons showing apparent treatment-specific clustering of mutations (*MXAN_2661, MXAN_5030, MXAN_7213*), we tested whether repeated mutations in the same operon were disproportionately concentrated within specific treatments, adopting the approach of Deatherage *et al.* 2017. For each operon, we excluded the first within-treatment mutation (since there was no *a priori* expectation regarding what treatment that mutation would occur in) and tested whether additional mutations were biased toward a particular treatment or treatment combination (*i.e.* the two treatments with prey) relative to random expectation using two-tailed Fisher’s exact tests, with resulting *p* values corrected for multiple testing with the Bonferroni method. *MXAN_2661* was mutated in six prey-adapted populations and one NP population, but this concentration in the prey-adapted populations was not significant (*p* = 0.242). The *MXAN_5030* operon was the second-most mutated operon overall and contains three genes predicted to encode constituents of an outer membrane HAE-1 efflux pump (*MXAN_5030-5032*). All eight populations adapted to the two prey environments were mutated at this operon but only one NP population was mutated at the operon (Fig. 4c), a concentration unexpected by chance (*p* = 0.036). The *MXAN_7213* operon also evolved in a non-random distribution across treatments, with mutations in this operon occurring in all four NP populations and one Bs population (Fig. 4c; *p* = 0.036).

As a final approach to examining treatment-specific genome evolution, we adapted a method previously developed for detecting biased gene expression across gene-ontology (GO) categories^58^, to test for non-random concentrations of evolved mutations in specific GO categories (as well as operons) for each of the three treatments (Supp. Fig. 5a). For most cases in which non-random concentration of mutations within a given GO category was detected, it was detected in only one treatment (14 cases; Supp. Fig. 5a). Only six GO categories showed significant mutation concentration in more than one treatment (four in two treatments and two in all three treatments). That almost all cases of significant GO-category mutation concentration occurred in just one treatment or two treatments suggests treatment-biased evolution at the GO-category level (Supp. Fig. 5a).

#### Test for effects of adaptation to prey on soft-agar swarming

*MXAN_5030, lonD* and *frz* operons were previously identified as targets of selection among populations adapted to prey-free CTT hard or soft agar in MyxoEE-3^22^. Here *lonD* was mutated only in three prey-adapted populations (Table 2), suggesting possible prey-biased mutation as was detected for *MXAN_5030*. In the previous study, as here, the *frz* operon was mutated in almost all replicate populations, but the *MXAN_5030* and *lonD* operons were preferentially mutated among populations adapted to soft agar. We therefore asked whether the suggested bias of mutation in these two operons toward prey-adapted populations in this study might be due to prey lawns having physical characteristics similar to prey-free soft agar. If this were the case, the prey-adapted populations might be expected to swarm faster on prey-free soft agar, just as they generally swarm faster in their home prey environments. However, this was not the case in our experiments (Supp. Fig. 6), suggesting that aspects of the Bs and Ec environments other than surface rigidity imposed selection on the *MXAN_5030* and *lonD* operons.

**Table 2:** Loci mutated only in two or more prey-adapted populations.

| Locus | Treatment | Mutation | (Predicted) Function |
| --- | --- | --- | --- |
| *MXAN_1365* | Ec | Inframe-insertion (Ec-1),<br>nonsense mutation (Ec-2) | Hypothetical protein* |
| *MXAN_3106*<br>(*kilC*) | Ec | Frameshift insertion (Ec-2),<br>missense mutation (Ec-3) | Secretin of the Kil system<sup>39</sup> |
| *MXAN_6755* | Ec | Missense mutations (Ec-1, Ec-4) | 4-vinyl reductase 4VR domain-<br>containing protein* |
| *MXAN_4699* | Bs | Missense mutation (Bs-3),<br>nonsense mutation (Bs-4) | Putative L-lysine 2,3-aminomutase* |
| *MXAN_5030* | Bs | Nonsense mutations (Bs-2, Bs-4) | Efflux transporter, HAE1<br>family, outer membrane efflux<br>protein* |
| *MXAN_6662* | Bs | Missense mutation (Bs-1),<br>frameshift deletion (Bs-3) | Branched chain amino acid<br>ABC transporter, ATP-binding<br>protein* |
| *MXAN_5204*<br>(*rpoD*) | Bs | Missense mutations (Bs-1, Bs-4) | RNA polymerase sigma factor<sup>83</sup> |
| *MXAN_5031* | Bs & Ec | Nonsense mutations (Bs-3, Ec-2) | Efflux transporter, HAE1<br>family, MFP subunit* |
| *MXAN_3993*<br>(*lonD*) | Bs & Ec | Frameshift deletion (Bs-2),<br>nonsense mutation (Ec-4),<br>missense mutation (Ec-3) | Lon protease<sup>84</sup> |

\*annotations based on UniProt and KEGG

### The lysine 2,3-aminomutase gene *MXAN_4699* (A271P) is necessary for increased predatory swarming by a clone from population Bs-3

Nine genes were mutated in two or three populations adapted in the presence of prey but not in any NP population (Table 2); seven of those were mutated in two replicate populations within the same prey treatment (Table 2, Fig. 4g). One of those genes, *MXAN_3106* (*kilC*), was previously shown to have a major role in *M. xanthus* predation^39^. Another, *MXAN_4699,* was mutated in two Bs populations that showed i) large increases in home-environment swarming, and ii) the overall largest coincidental increases in swarming across our panel of foreign prey (Supp. Fig. 1d, Supp. Fig. 2c). Both of the *MXAN_4699* mutations in these populations – a missense mutation (A271P) in Bs-3 and a nonsense mutation (Q466*) in Bs-4 (Supp. Table 4) – went to fixation. Due to these patterns, we targeted *MXAN_4699* and the mutation of this gene in population Bs-3 for initial molecular-genetic analysis. Specifically, we sought to test for a functional role of the Bs-3 *MXAN_4699* mutation in the evolved predatory swarming phenotypes of this population, which exhibited the largest proportional increase in home-prey swarming of any of the 12 MyxoEE-3 populations examined here. *MXAN_4699* is predicted to encode a 456-aa long lysine-2,3-aminomutase (KAM), which catalyses the conversion of lysine to β-lysine^59,60^.

We first swapped the mutated Bs-3 *MXAN_4699* allele (A271P) with its ancestral variant in a clone (Bs-3.2) isolated from population Bs-3 that exhibits a home-environment swarming phenotype similar to that of the whole Bs-3 population (Fig. 5b). However, no significant effect of this mutation reversion on either predatory swarming or competitive performance against the ancestor in the *B. subtilis* home environment of Bs-3 was observed (Fig. 5b, Supp. Fig. 7b). Despite these null results, we reasoned that this mutation may have caused adaptive phenotypic effects in the genetic background in which it first occurred that are masked by effects of one or more mutations that occurred later in the Bs-3 lineage.

**Figure 5.**
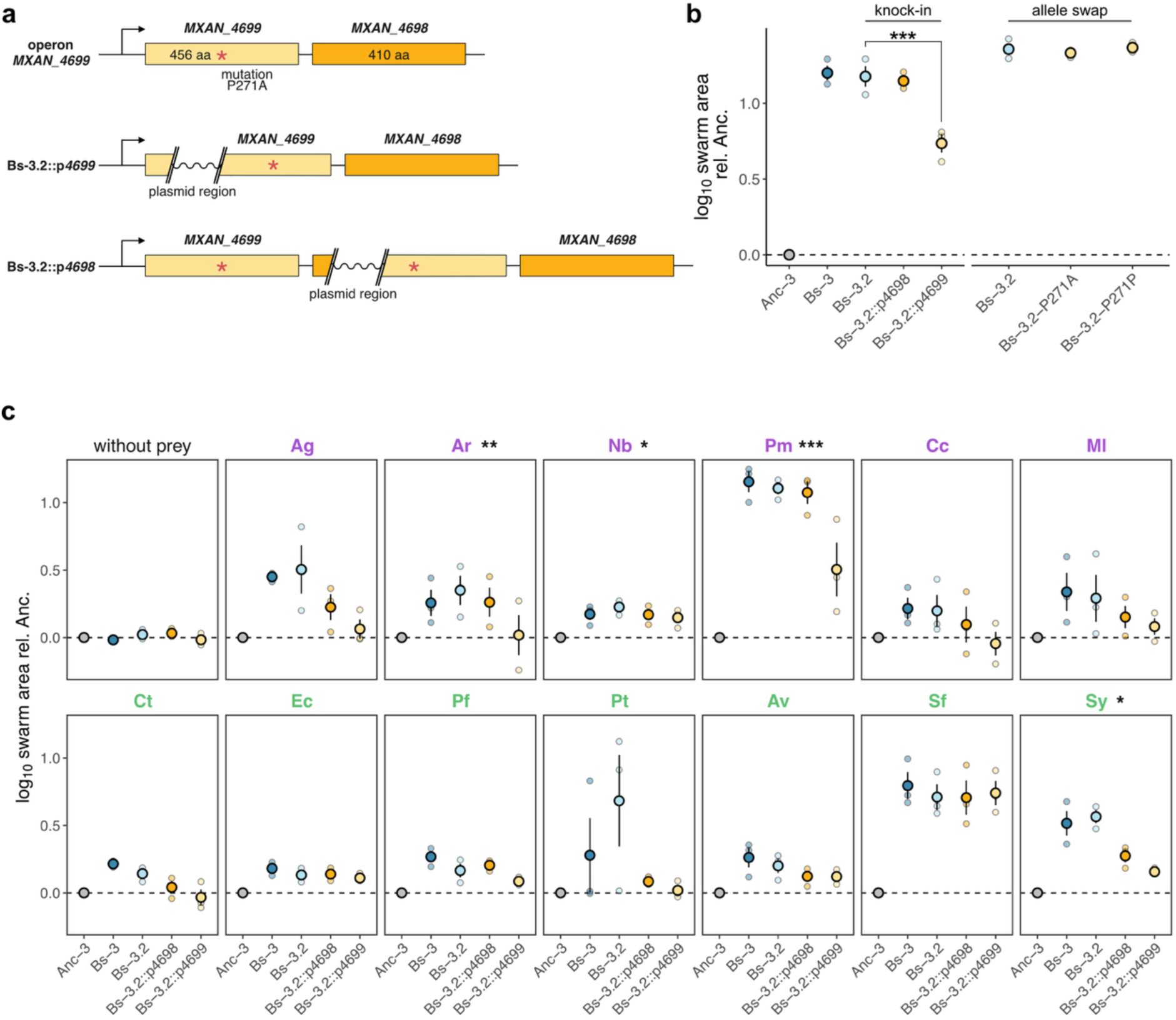
Disruption of *MXAN_4699* strongly reduces evolved predatory swarming. **a)** Schematic of the *MXAN_4699/4698* operon and disruption insertions in both genes. Both *MXAN_4699* and *MXAN_4698* are annotated as putative lysine-2,3-aminomutases. *MXAN_4699* is mutated in populations Bs-3 and Bs-4, which both showed increased swarming on *B. subtilis*. The red asterisk shows the approximate position of the mutation in Bs-3. The plasmid that was used for creating the insertion knock-outs consists of a pCR-Blunt backbone and a fragment which aligns to the beginning of the target gene. Once the plasmid crosses in, it separates the beginning of the gene and its promoter from the remainder of the gene, rendering the gene non-functional. **b)** Swarm areas on *B. subtilis* lawns relative to the clonal ancestor of Bs-3 (Anc-3): from left to right Anc-3, the evolved Bs-3 population, a clone isolated from the Bs-3 population and used for genetic manipulations (Bs-3.2), two plasmid-disruption mutants of Bs-3.2 (Bs-3.2::p4698 and Bs-3.2::p4699), a derivative of the allele-exchange process that retained the ancestral allele at *MXAN_4699* (Bs-3.2-P271A, see Methods), and a derivative of the allele-exchange process carrying the evolved allele at *MXAN_4699* (Bs-3.2-P271P). The plasmid disruption in Bs-3.2 (Bs-3.2::p4699) renders both *MXAN_4699* and *MXAN_4698* non-functional. For the allele swap, a clone with the reversion to the ancestral allele (Bs-3.2-P271A) and a clone which retained the mutated allele (Bs-3.2-P271P) were assayed in parallel to control for any effects stemming from the genetic manipulation process. **c)** Swarm areas on lawns of diverse prey, relative to the ancestral clone Anc-3. The plot on the top left shows swarm areas on CTT hard agar without prey. For the remaining panels, the top and bottom rows show results from lawns of Gram-positive and Gram-negative prey, respectively. Error bars show the SEM for three biological replicates. Asterisks next to the prey species indicate a significant difference between the unmanipulated clone Bs-3.2 and the knock-in clone Bs-3.2::p4699 (* *p* < 0.05, ** *p* < 0.01, *** p <0.001).

We therefore tested whether having a functional copy of the *MXAN_4699* gene as a whole is important for Bs-3.2 evolved phenotypes. We additionally tested for any role of *MXAN_4698,* which lies downstream from *MXAN_4699* in the same operon (Fig. 5a) and is predicted to be a KAM that shares 23% amino-acid identity with *MXAN_4699*. We constructed two mutants: one with an insertion at the beginning of *MXAN_4698* and another with an insertion at the start of *MXAN_4699*. These long inserts are expected to knock out the respective gene’s function by preventing transcription of the majority of the gene. Importantly, because of the polycistronic nature of the operon, we expect the disruption of *MXAN_4699* to render both genes non-functional (Fig. 5a).

Disruption of *MXAN_4699,* resulting in non-functional *MXAN_4699* and *MXAN_4698*, greatly reduced the evolutionarily increased swarming of Bs-3.2 on *B. subtilis,* although not fully back to the ancestral state (Fig. 5b). However, disruption of *MXAN_4698* alone did not cause any reduction, indicating that only *MXAN_4699* with the mutation (A271P) is necessary for full expression of the evolved predatory swarming phenotype of Bs-3.2.

We additionally asked whether having a functional copy of *MXAN_4699* (A271P) with or without *MXAN_4698* is necessary for the coincidentally increased swarming by population Bs-3 observed on foreign prey (Fig. 5c). On a subset of the foreign prey, we observed differences in swarm size between the whole evolved population Bs-3 and the isolated clone used as the parent of our mutants (Fig. 5c), despite their similar phenotype on *B. subtilis* (Fig. 5b). Given these differences, we used Bs-3.2 as the reference point for assessing the disruption mutants.

The impact of knocking out both *MXAN_4699* and *MXAN_4698* or *MXAN_4698* alone on predatory swarming varied depending on the prey species. Disruption of *MXAN_4699* and *MXAN_4698* clearly reduced swarming on most foreign prey species (Fig. 5c), for example on *A. ramosus, N. bataviensis, P. megaterium,* and *S. yanoikuyae* (*p_Ar_*= 0.001, *p_Nb_*= 0.013, *p_Pm_*< 0.001, *p_Sy_*= 0.018). On some of those same foreign prey, knocking out only *MXAN_4698* had no major effect (e.g. *A. ramosus*, *P. megaterium*), implicating a role for *MXAN_4699* (A271P) alone. On a few species however, knocking out *MXAN_4698* also had a clear or suggestive negative effect on swarming (e.g. *A. globiformis, C. testosteronii, P. tolaasi and S. yanoikuyae*), either an effect similar in degree to the *MXAN_4699-MXAN_4698* knock-out or a lesser effect. For some prey, neither *MXAN_4699-MXAN_4698* nor *MXAN_4698* knock-outs appear to have reduced swarming compared to the original clone (*S. fredii, E. coli*).

Collectively, the results of our genetic manipulations identify a gene necessary for full expression of the strikingly large evolutionary increase in the swarming ability of an evolved clone adapted to preying on *B. subtilis,* but without finding a functional effect of the evolved mutation in that gene in the genetic background of the terminal clone. Future work is necessary to investigate potential effects of that mutation in the genetic background in which it first appeared.

## Discussion

How evolution in one environment indirectly shapes performance and phenotypes in others through pleiotropy, hitchhiking and relaxed selection is a central question in evolutionary theory^1–3,6^. While antagonistic pleiotropy has dominated this theme^5,27,61^, evolution experiments with microbes have shown that indirect effects of adapting to one set of abiotic conditions can be positive on average in other conditions^26,28^, resulting in positive specialization (Fig. 1a). Addressing biotic interactions, Stewart *et al.* (2022) found that transposon-insertion mutations introduced into the protist *Dictyostelium discoideum* that improved predatory fitness on one prey type generally had little effect on fitness with other prey (neutral specialization, Fig. 1a) suggesting that adaptive predatory specialization might often proceed without general tradeoffs.

Our results with a long-term evolution experiment support this hypothesis and go further to suggest that predation-related adaptation to one prey environment fueled by spontaneous mutation may often increase average fitness across other prey environments, generating positive specialization. Evolving populations of the generalist bacterial predator *M. xanthus* adapting to non-evolving lawns of a single bacterial prey species increased in swarming rate on their home prey while also increasing to a lesser degree in their average swarming rates across diverse foreign prey (Fig. 3a).

Yet the observed trend of positive specialization masks much idiosyncrasy in the magnitude and sign of indirect effects, which caused striking radiation of predation profiles through combined effects of selection and chance. The character of indirect evolution varied with respect to several factors: i) MyxoEE-3 prey treatments – reflecting historical selective effects of prey type (Fig. 3a, d), ii) replicate populations within treatments – reflecting chance effects on mutational pathways (Fig. 3c), iii) foreign-prey identity – reflecting ecological effects of prey environments not encountered during selection (Fig. 3c), and iv) interactions among the preceding three factors. Among the historical selective effects, populations adapted to *B. subtilis* showed a greater average increase on foreign prey (Fig. 3a) and greater chance diversification among replicate populations across foreign prey than populations adapted to *E. coli* (Fig. 3d, Supp. Fig. 3).

At the genomic level, populations tended to evolve more similarly within treatments than between them (Fig. 4a-f). Such parallel evolution was distributed across many loci, indicating that prey-specific adaptation draws on a broad genetic architecture rather than few large-effect genes. Treatment-specific genomic parallelism was strongest in the *E. coli-*adapted populations (Fig. 4b, e), an outcome associated with lower phenotypic diversification (Supp. Fig. 1, Fig. 3c). The greater phenotypic and genomic diversification among replicates observed under *B. subtilis* selection illustrates how ecological context can modulate the balance between determinism and chance in evolution^9,49,51,62,63^.

Environmental effects on the relative evolutionary roles of selection and chance can be mediated by effects on growth rate, population size^63^ and/or by environment-specific differences in the structure of adaptive landscapes^9,62,64^. Both hypotheses might apply to our results given that swarm expansion by our ancestor is slower on *B. subtilis* than on *E. coli* (Fig. 1a). This ancestral performance differential may have been associated with more adaptive-mutation pathways being accessible in the Bs environment, and implies that the Bs populations underwent fewer generations per cycle than the Ec populations early during MyxoEE-3.

### Nested parallel evolution

We observed nested levels of parallel evolution, with some genetic targets of selection mediating adaptation generically to both prey and non-prey MyxoEE-3 environments (*e.g.* the *frz* operon that controls cell-reversal frequency^49,56^) and others that appear to have (or may have) mediated prey-specific adaptation (Fig. 4c, f; Supp. Fig. 5a; Table 2). This structure reflects joint effects of selective features shared across treatments (*e.g.* the need to be near the leading front of expanding swarms to be propagated across cycles) and prey-specific selective factors that remain to be identified and distinguished. The observed cases of prey-specific parallelism enrich future investigation of the genetic basis of predatory fitness in *M. xanthus.* Most of the identified candidate loci have not previously been associated with predatory fitness, suggesting that the genetic architecture underlying *M. xanthus* predatory fitness is more complex than previously understood. Two of these candidate loci have been experimentally connected to predation to date – *MXAN_3106* (*kilC*) from previous studies^39^ and *MXAN_4699* from work described here, suggesting that other loci in our list will be shown to also shape predation phenotypes in future work.

The lysine-2,3-aminomutase gene *MXAN_4699* was found to be required for full expression of evolutionarily increased swarming by a Bs-3 clone on *B. subtilis* lawns (Fig. 5b). Surprisingly however, despite the requirement for a functional copy of this gene as a whole, the mutation which first alerted us to *MXAN_4699* was not required for the evolved clone’s increased-swarming phenotype, either on *B. subtilis* or on foreign prey (Fig. 5b, Supp. Fig. 7c). This result suggests that the mutation may have increased fitness in the genotype in which it first occurred but that its phenotypic effects became masked by effects of mutations that accumulated later in the clone’s lineage.

A given set of evolved mutations present in an evolved population often had very different effects on predatory swarming across different prey species (Fig. 3c). For the *MXAN_4699* disruption, resulting in non-functional *MXAN_4699* and *MXAN_4698,* we see the same outcome of very different mutation effects in different prey environments, *i.e.* a strong genotype x environment (G x E) interaction. Disruption of *MXAN_4699* in a clone from population Bs-3 clearly or suggestively reduced evolved swarming gains on most tested prey species (e.g. *P. megaterium*) while having little or no effect on others (e.g. *S. fredii)* (Fig. 5). Thus, having a functional copy of the mutated gene contributes to evolutionarily increased swarming in some prey environments but not others.

### Indirect evolution and microbial life cycles

Evolution experiments with diverse microbes have demonstrated the importance of indirect effects in the evolution of growth rates, metabolic functions and stress resistance in largely abiotic contexts^3,10,26^. Studies with *M. xanthus* have expanded this approach to behavioral and interaction phenotypes, including developmental, motility, social, predatory and pathogen-resistance phenotypes^9,11,12,22,35,51^, themes central to animal and plant evolution. For example, adaptation to the same *B. subtilis* and *E. coli* MyxoEE-3 selective regimes examined in this study was found to indirectly alter morphological and sporulation phenotypes during multicellular fruiting body development differently than evolution without prey (the NP treatment here)^22^. Moreover, the two prey species exerted different indirect effects on development, mirroring the home-prey-specific differences in indirect predation effects observed here. Our present study expands study of latent-phenotype evolution^9^ to behavioral radiation across diverse prey environments. Collectively, these studies show that indirect evolution is a major force in the evolution of myxobacterial life cycles, sociality and inter-species interactions and point to a complex network of pleiotropic interactions across traits. Elucidating the character of such networks is important for understanding pathways by which and constraints on how microbial life cycles and predation profiles evolve. In turn, how predation profiles evolve directly and indirectly has implications for the use of experimental evolution to enhance microbial predators as biocontrol agents in agriculture^65^ and medicine^66,67^.

### Natural diversity

The predatory radiation observed here under simple, single-prey selection provides insight into how the observed diversity of predatory profiles in natural *M. xanthus* populations^32,33^ can arise. In spatially structured soils, local prey communities vary across both space and time, exposing predators to distinct biotic selective regimes. Such variation in prey environments will shape differentiation of predator subpopulations not only by shifting the genetic targets of selection and selection strength across microhabitats, but also by influencing the role of chance in predator diversification, whether by shifting the structure of adaptive landscapes or impacting predator population size.

Ecologically, predatory specializations are expected to promote co-existence of diversified predator lineages. Functionally distinct predators can partition prey species and spatial niches, reducing direct competition^13^. In turn, differential predation pressures from diversified predators shape community diversity^68^ and can promote maintenance of prey diversity^69^.

Adaptation by predators to even single-prey environments can generate a rich array of indirect outcomes shaped by pleiotropic effects of mutations accumulated during adaptation and ecological specificities of predator-prey interactions. Such coincidental effects can be a creative force in predator-prey evolution, generating novel functional diversity even under relatively constant and simple selective conditions. Our results point to how biotic context, pleiotropy and chance can interact to shape the emergence of specialized and diverse predator profiles.

## Methods

### MyxoEE-3

In this study, we used a subset of evolved populations from a broader evolution experiment which was performed from 2001-2003 and in 2020 was named MyxoEE-3^11,22,45,46,51^, where ‘MyxoEE’ stands for “Myxobacteria Evolution Experiment” and “3” refers to the temporal rank position considering the first MyxoEE-3 publication relative to those from other MyxoEEs (www.myxoee.org/). The 12 MyxoEE-3 populations examined here were founded by four distinct sub-clones of *M. xanthus* strain GJV1 (here referred to as “Anc” for ancestor), which were used to establish four replicate populations in each of three evolutionary treatments that differed in their selective environment (Supp. Table 3). All three treatments shared a physical base of CTT 1.5% agar [8 mM MgSO_4_, 10 mM Tris-HCl pH 8.0, 1% casitone, 1 mM K_3_PO_4_, 1.5% *Select* agar; pH = 7.6]; one had no prey added (No-Prey = NP), one had lawns of *B. subtilis* (Bs) and one had lawns of *E. coli* (Ec) (Fig. 1). The treatments referred to as ‘NP’, ‘Bs’ and ‘Ec’ here for simplicity were referred to as ‘CTT HA, ‘CTT HA *B. subtilis*’, ‘CTT HA *E. coli*’ in Rendueles *et al.* 2015, which was the first publication to examine populations from multiple MyxoEE-3 treatments. The populations examined here evolved for 40 cycles, corresponding to the terminal cycle of the MyxoEE-3.

### Swarming Assay

As noted in the Introduction, the rate at which *M. xanthus* swarms through lawns of prey, while not a complete measure of predatory fitness, has been shown to correlate with several other measures of predatory performance^44^. Most fundamentally, it determines the rate at which an expanding *M. xanthus* colony encounters new prey that predator cells can potentially kill and consume. Here we adopt this trait as our primary measure of performance on both home and foreign because it is clearly important for predatory fitness in our experimental regime overall and could be readily assayed across a wide range of foreign prey types.

All swarming assays were conducted on 9-cm diameter Petri dishes containing 30 ml of CTT 1.5% agar. All plates were prepared one day before the experiment; the autoclaved agar was added to plates and left to solidify under laminar airflow for about 45 min uncapped before being covered. Excluding *M. xanthus*, all bacterial species were inoculated into CTT liquid media [8 mM MgSO_4_, 10 mM Tris-HCl pH 8.0, 1% casitone, 1 mM K_3_PO_4_, pH = 7.6] the same day plates were prepared and then grown to stationary phase overnight. From the stationary phase culture, 129 µl were spread uniformly over the entire surface of a CTT-agar plate and then allowed to dry uncapped for ∼1 hr under laminar airflow. Cultures of *M. xanthus* cells were grown to mid-log phase and then pelleted and resuspended in CTT liquid to ∼5×10^9^ cells/mL. 10 µL of resuspended culture were spotted in the centre of the prepared prey plates and allowed to dry open under laminar airflow for ∼30 min. Once inocula had dried, plates were incubated upside down at 32 °C, 90% rH. Perimeters of *M. xanthus* swarms were manually marked every 24 h for up to seven days. Volumes and cell densities were calculated to replicate the conditions used in the MyxoEE-3.

### Competition Assay

Because most strains of *M. xanthus,* including many of those examined in this study, are not easily dispersed after swarming on an agar surface, dispersion and dilution plating to estimate population sizes of strains competing in co-culture was not optimal for this study. We therefore used an alternative approach that allowed us to indirectly estimate the population size of a marked variant of our ancestor after it had competed with either the unmarked ancestor or unmarked evolved populations under specified conditions. By doing so, we could quantify a readout of the competitiveness of evolved populations relative to their ancestor as a proxy for a more direct measure of relative fitness.

In short, post-competition population size of the marked ancestor was interpreted as inversely reflecting the competitiveness of its unmarked competitors, with values from competitions with the unmarked ancestor serving as the ancestral baseline. Direct estimation of relative fitness was not possible because our competition assay did not allow estimation of the population size of the unmarked competitor. A conceptually similar competition assay was utilized in a previous study of other MyxoEE-3 populations^49^, except the output of that assay was simply scoring the presence or absence of the unmarked ancestor from population samples taken from the outer edge of competition swarms, with absence of the ancestor reflecting clear fitness superiority of a competing strain. Our current assay improves on the earlier assay by allowing estimates of evolved competitiveness along a continuous scale.

Competition assays were carried out using subclones of the strain GJV2, a rifampicin-resistant mutant derivative of the GJV1 strain from which MyxoEE-3 ancestral sub-clones were isolated (see Methods: MyxoEE-3). GJV2 resistance to rifampicin was previously shown to result from a single base-pair mutation in the gene *rpoB*. Here, GJV1 and GJV2 are also referred to as Anc and Anc^Rif^, respectively. Anc^Rif^ population size after competition in initially 1:1 mixes with evolved populations vs with Anc was used as a measure of evolutionary change in the competitiveness of evolved populations versus their ancestor Anc^Rif^. Competitiveness was calculated according to the equation shown in Supp. Fig. 1b, such that positive values reflect increased competitiveness compared to the ancestor. Our overall competition assays were conducted in two stages: i) the competition itself (Stage 1) and ii) estimation of post-competition sizes of Anc^Rif^ populations (Stage 2).

*M. xanthus* swarm size after a defined period has previously been shown to correlate with initial inoculum size^70^. To allow estimation of Anc^Rif^ population sizes at the end of Stage 1, standard curves based on the area of four-day swarms were quantified in parallel with each competition assay for Anc^Rif^ clones from pure-culture populations spotted at several known starting cell densities (5×10^9^ cells/mL; 5×10^8^ cells/mL; 5×10^7^ cells/mL; 5×10^6^ cells/mL) on CTT agar plates containing rifampicin [5 µg/mL] (Supp. Fig. 1c).

### Stage 1: Competition

Petri dishes of 6 cm diameter were prepared two days before the experiment with 10 ml of CTT agar and left to dry under laminar airflow. Populations of *E. coli* and *B. subtilis* cells were grown independently to stationary phase overnight in CTT liquid. From these cultures, 71 µL were spotted and spread over the surface of the previously prepared plates and left to dry under laminar airflow for about 30 min. Consistent with all assays, exponentially growing cultures of *M. xanthus* were pelleted and resuspended in CTT liquid to a density ∼5×10^9^ cells/mL. Resuspended pellets of all evolved populations and ancestors were mixed at a 1:1 ratio with the rifampicin-resistant ancestral subclones of GJV2 (Anc^Rif^). Of the obtained mixes, 10 µL were spotted at the centre of plates with the MyxoEE-3 selective environment in which the respective population had evolved. Control mixes consisting of GJV1 and GJV2 cells (Anc vs Anc^Rif^) were spotted to quantify the performance of the ancestral subclones in each selective environment. *M. xanthus* spots were left to dry for about 30 min and then incubated upside down at 32 °C with 90% rH.

### Stage 2: Estimation of post-competition Anc^Rif^ population size

After five days of incubation, *M. xanthus* swarms were scraped from the agar surface and resuspended in 1 mL of CTT liquid. The resuspension of cells from the collected swarms was achieved mechanically by vigorously vortexing tubes containing glass beads (three 3 mm beads for each 1,5 mL tube) for 30 sec and subsequently pipetting up and down 500 µL of the resuspension for 30 sec. 50 µL of the resuspended cells were spotted on new Petri dishes (9 cm diameter) containing 20 ml CTT 1.5% agar with rifampicin (5 µg/mL) and left to dry under laminar airflow for about 30 min. Once dried, plates were incubated at 32°C with 90% rH for four days, after which the *M. xanthus* swarm areas were marked and quantified.

### Growth curves

Growth curves were measured as change in OD_595_ over time with a *Tecan infinite M200 Pro* plate reader. After growing the evolved populations and their ancestors to mid-exponential phase in CTT liquid, the cultures were diluted into fresh CTT liquid to a starting OD_595_ of ∼0.1. 300 µL of each diluted culture were placed in a Greiner 96-well flat-bottom plate. To minimise any influence of well location, all samples were measured in two randomly assigned wells in the same plate, with random assignments performed independently for each assay replicate. Each replicate was run for 48h, taking five reads per well every 30 minutes. Between measurements, the machine was set to keep the temperature at 32°C and shake the plate. The raw data generated by the plate reader in an Excel file was processed using the R package *tread*^71^. For further analysis, the average of the five reads for each well was taken and further averaged across the two technical replicates (measurements of the same strain in separate wells).

### Image acquisition and swarm quantification

All swarms were manually traced with a marker directly on the back of the plates, which were then photographed with a Nikon Coolpix S10 camera with the same imaging conditions across replicates and experiments. The areas of the previously traced swarms were measured in *Fiji*^72^ (version 2.3.0/1.53q) by manually creating a *freehand selection* and then measuring the obtained selection area expressed in px^2^. In the case of multi-day measurements, a three-colour code was used to help distinguish multiple individual traces drawn on the same plate over multiple days.

### DNA extraction and quality control

Evolved populations were inoculated from freezer stocks and grown directly in CTT liquid containing gentamicin (1 µg/mL, to prevent contamination) until cultures reached OD_595_ 0.4-0.8. Then, cells were centrifuged at 12000 rpm for 5 min, the supernatant discarded, and the obtained pellet frozen and kept at -80 °C until pellets from all evolved populations were collected. Frozen pellets were thawed at room temperature, and the *QIAGEN* Genomic DNA Buffer Set was used to extract total genomic DNA according to the manufacturer’s protocol. However, given the high cohesiveness of some *M. xanthus* populations, we extended the incubation time in the lysis buffer from 30 min to 45 min at 37 °C. The *QIAGEN* Genomic-tip 20/G kit was used to perform the washing steps, leading to the purification of the extracted DNA. In some cases, due to the tendency of the lysate debris to clog columns, the buffer required for the washing steps needed to be forced through by manually increasing the pressure. The obtained genomic DNA was resuspended in EB buffer, the quality was assessed with a NanoDrop spectrophotometer, and DNA concentration was measured with a Qubit fluorometer.

### Genome sequencing and mutation calls

Library preparation and sequencing was performed at Oxford Genomics, Oxford (UK). In brief, DNA quantification was conducted employing either Qubit (Invitrogen) or PicoGreen (Thermo Fisher) on the FLUOstar OPTIMA plate reader (BMG Labtech). The input material was normalized to 500 ng before undergoing fragmentation and library preparation. Fragmentation was achieved through mechanical shearing, targeting an average size of 350 bp, utilizing a MultiFunctional Bioprocessor (EpiSonic; amplitude 40, process time 3 min 20 sec, pulse on/off 20 sec). Library preparation involved the NEBNext Ultra DNA library prep kit for Illumina (New England Biolabs) and standard Illumina multiplexing adapters, with slight modifications to the manufacturer’s protocol. PCR amplification (10 cycles for isolate samples and 14 cycles for meta samples) occurred on a Tetrad (Bio-Rad) using in-house unique dual indexing primers^73^. Post-PCR purification was carried out using Agencourt Ampure XP (Beckman Coulter; ratio 1:0.75). Individual libraries were normalized using PicoGreen (Thermo Fisher) or Quantifluor (Promega), then pooled accordingly. The size profile of the pooled library was assessed on the 2200 or 4200 TapeStation. The final pooled library was quantified using Qubit (Invitrogen) and diluted to approximately 10 nM for storage. The 10 nM library underwent denaturation and further dilution before loading onto the sequencer. Paired-end sequencing was executed on a NovaSeq6000 platform (Illumina, NovaSeq 6000 S2/S4 reagent kit v1.5, 300 cycles), thereby generating 150bp-reads.

Raw read quantity and quality was assessed with *FastQC* v0.11.4^74^ and trimming, and adaptor-removal were performed in paired-end mode (PE) using *trimmomatic* v0.35^75^ with the following parameters (where ${adapter} refers to concatenated file jointly containing the programme’s TruSeq and Nextera adapters):

> ILLUMINACLIP:${adapter}:2:25:10 CROP:149 LEADING:30 TRAILING:28 SLIDINGWINDOW:4:28 MINLEN:36

To assess mutational frequencies in whole population samples, we first removed duplicates using *picard-tools* v2.21.3^76^ from all trimmed reads, and remapped these deduplicated reads using *breseq* v0.35.5^77^ in population mode (-p) against the reference genome of *M. xanthus* DK1622 (RefSeq: 13-FEB-2022; GenBank Accession NC_008095) with parameters suited for population mapping (*-p*), and considering a mapping quality filter of 25 (*--minimum-mapping-quality 25*), as well as a minimum mapping requirements min. 10 mapping reads that support either variant or ancestral base, respectively (*--consensus-minimum-total-coverage 10 --consensus-minimum-variant-coverage-each-strand 10*).

Mutations and their respective within-population-frequencies were annotated for all whole population samples using *breseq*’s internal summary function “*gdtools* ANNOTATE” relative to DK1622, and count statistics were generated using “*gdtools* COUNT -b”. We annotated the effects of all mutations in coding and intergenic regions and inferred dN over dS ratios from the inferred COUNT data for all fixed (100%) single nucleotide variants (SNV) in coding regions bytakinbg into account the observed sites at risk to produce nonsynonymous and synonymous sites, respectively, for each evolved population.

### Allele swap and construction of insertion knock-in mutants

For all genetic manipulations, clones were first isolated from the original evolved population Bs-3. The swarm area on *B. subtilis* was measured for each clone to ensure that they show the same phenotype as the evolved population.

For constructing knock-in mutations in *MXAN_4698* and *MXAN_4699*, plasmids (p4698 and p4699, Supp. Table 1) were constructed with the pCR-Blunt vector (Zero Blunt PCR Cloning Kit, *Invitrogen*), which contains a kanamycin resistance gene. To create plasmid p4699 and p4698, a fragment was amplified with primer sets GV919 and GV921 or GV1147 and GV1149, respectively (Supp. Table 1). The fragments were then ligated with the linearised pCR-Blunt backbone using T4 ligase. Clones of Bs-3 were transformed with the resulting plasmid through electroporation and the potential mutants (Bs-3.2::p4699 and Bs-3.2::p4698) were subsequently selected with kanamycin (40 μg/ml) for uptake of the plasmid. For all subsequent experiments, the knock-in clones were cultured with kanamycin to maintain the plasmid, except when grown in liquid before starting an experiment which involved prey or kanamycin-sensitive *M. xanthus* strains.

An allele exchange was performed to replace the mutation in the evolved clone Bs-3.2 with the ancestral *MXAN_4699* allele. The plasmid pCR-4699 was constructed by ligation with a PCR fragment of the ancestral *MXAN_4699* (amplified with primers GV1147 and GV1148), which extends ∼500bp on both sides of the mutation in population Bs-3. The PCR fragment was then extracted from pCR-4699 and cloned into the pBJ vector^78^. The resulting plasmid pBJ-4699 contains both a kanamycin resistance gene and a *galK* gene. Bs-3.2 was transformed with pBJ-4699 and selected for uptake of the plasmid with kanamycin. Successful transformants were then grown in CTT liquid without kanamycin and counter-selected with galactose for the loss of the plasmid. By losing the plasmid, the mutant has roughly an equal chance of retaining either the original mutation or wildtype allele. The resulting clones can be verified by Sanger sequencing for successful allele exchange (Bs-3.2-P271A). Clones which retained the original mutation were used as controls in experiments (Bs-3.2-P271P).

### Data analysis

All experiments were performed in three temporally independent (*i.e.* initiated on three different days with independently prepared media) replicate blocks. Every replicate block included the ancestral sub-clones as internal control used to determine degrees of evolutionary change and divergence for the assessed trait in all experiments. The software R v4.5.0 was used for all statistical analyses and generating graphs^79^. All data sources include the details of the specific statistical analyses used.

#### Enrichment analysis

Non-random concentration of mutations within gene-ontology (GO) categories and operons was determined with *FUNAGE-Pro v2*^58^, a comprehensive web server dedicated to the functional analysis of prokaryotic genomes. Statistical significance was evaluated using the hypergeometric distribution, modeling the probability of drawing the observed number of mutated genes belonging to a given functional class out of the total set of mutated genes when sampling without replacement from the complete genome. Raw *p* values were corrected with Benjamini-Hochberg for multiple testing across all tested categories. *FUNAGE-Pro* draws on a curated database containing multiple functional classifications of genes derived from all publicly available complete bacterial genomes, supporting common annotation systems such as GO, COG, KEGG, InterPro, PFAM, eggnog, and predicted operon classes. The mutation lists, analyzed for each treatment separately, were submitted as a single gene set and queried against the *Myxococcus xanthus* DK1622 reference genome (assembly ASM1268v1) to retrieve the corresponding functional annotations. However, only GO and operon classes were considered for this analysis since 40% of genes were classified as poorly characterized according to the COG system (Supp. Fig. 5b). Assignment of genes to COG classes was done using the list of COGs from La Fortezza *et al.* 2025. Operons were predicted from the reference genome of DK1622 using the web service *Operon-Mapper*^81^.

#### Interaction networks

For all possible pairwise population comparisons, the number of mutated genes or operons shared by the two populations in each pair (irrespective of the actual mutation) was calculated. The resulting data were used to construct an adjacency matrix that was plotted as an interaction network using the R function *qgraph*^82^, where the opacity and thickness of the connecting lines represented the number of shared mutated operons or genes. Operon names were assigned based on the first gene present in each specific operon on the sense or antisense strand.

#### Soerensen-Dice similarity index

The SD coefficient is calculated as the ratio of twice the number of shared mutated operons or genes and the number of mutated operons or genes unique to each population^57^. For within-treatment similarity the coefficient was calculated for each pair of populations within a given treatment and averaged. For across-treatment similarity the coefficient was calculated for all possible pairwise combinations of populations across treatments and averaged. This procedure resulted in 18 and 48 pairwise combinations total for within- and between-treatment pairs, respectively. Significance of the difference between within-treatment and across-treatment similarity was tested for by randomisation of treatment assignment of individual populations (adapted from Deatherage *et al.* 2017). The association between populations and their respective set of mutations was kept constant, but the populations were randomly assigned to the three treatments (NP, Ec, and Bs, always four replicate populations per treatment). For each random assignment of treatment labels, the mean similarity within and across treatments was calculated and the difference compared to the actual observed difference. Given 12 populations split into three treatments, 34,650 unique assignments of treatment labels are possible. Out of all possible assignments, the difference between within- vs across-treatment SD values was larger than or equal to the observed difference for the one true set of treatment assignments in only 12 cases for operons (*p* = 0.00035), and only 114 cases for genes (*p* = 0.00329). The SD coefficient considers only the presence or absence of shared mutated loci, ignoring the specific within-population frequencies of individual mutation.

For gene- and operon-level analyses (data for Fig. 4), intergenic mutations in CRISPR arrays were excluded, since they are not part of a gene or operon. A total of 20 mutations were excluded, all of them at low frequency (5-16%) in their respective populations.

### Mutational probabilities

Mutational probability analyses were performed to evaluate whether the recurrence of mutations within specific genes or operons across replicate populations and treatments exceeded that expected by chance. Because the first occurrence of a mutation at a locus was not considered unexpected *a priori*, probabilities were calculated conditional on one observed hit. Thus, we considered the likelihood that additional mutations occurred in the remaining 11 populations.

#### Genome-wide recurrence across populations

The probability that any given gene or operon would be mutated in multiple populations by chance was estimated considering the total number of mutations (240) across the twelve experimental populations with a binomial model. Mutations were assumed to occur independently, to be randomly distributed among loci, and to be equally likely to occur in any given locus of either type (gene or operon). For genes (7454 in DK1622) and operons (4293 in DK1622), the probability of being mutated in two or more populations is 0.029 and 0.050, respectively (excluding the first occurrence of a mutation as described above). The probability of being mutated in three or more populations is 0.0004 for genes and 0.0011 for operons, respectively.

## Acknowledgements

We thank Michael Manhart and Macarena Toll-Riera for their helpful comments on the manuscript and the Evolutionary Biology group at ETH Zurich for all the stimulating and inspiring discussions on this work.

## Author contributions

GJV, ML and SS designed experiments and wrote the manuscript. ML, SS, SF and YTNY performed experiments. SW and ML analysed the sequencing data, ML and SS created figures and performed statistical analysis for all datasets. GJV, ML, SW, YTNY, and SS revised the manuscript.

