## Supplemental Figures and Tables for "Predator adaptation to single prey species yields positive specialization and multiple forms of diversification"

### Supplementary Information

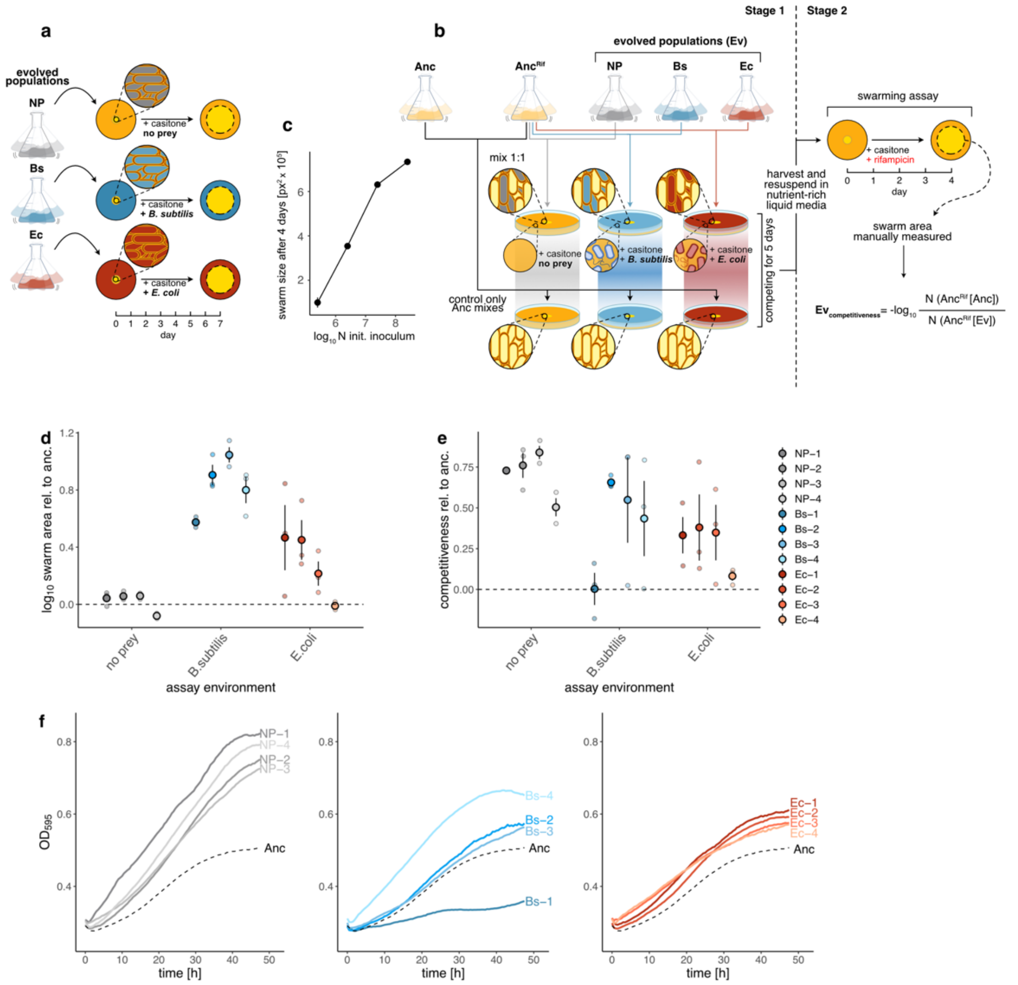


$$Ev_{competitiveness} = -\log_{10} \frac{N\,(\text{Anc}^{\text{Rif}}\,[\text{Anc}])}{N\,(\text{Anc}^{\text{Rif}}\,[\text{Ev}])}$$

**Supp. Figure 1. Swarm expansion and competition assays. a)** Illustration of the swarm-area assay featuring evolved populations assayed in their home MyxoEE-3 environment, as used to generate the data shown in Fig. 2b. **b**) Diagram illustrating competition assays used to generate the data shown in Figs. 2c and 2b. The overall assay was composed of two phases, the competition phase (Stage 1) and a second phase in which the post-competition sizes of the marked-ancestor populations were estimated (Stage 2). (See Methods for greater detail.) In Stage 1, freezer-stock samples of the terminal evolved populations (Ev) and the rifampicin-resistant ancestral clones ($\text{Anc}^{\text{Rif}}$) were grown in CTT liquid medium to mid-exponential phase. Ev and $\text{Anc}^{\text{Rif}}$ cells were mixed 1:1 and plated at the center of NP, Ec, and Bs plates. In parallel, the original rifampicin-sensitive ancestral clones (Anc) were similarly mixed with $\text{Anc}^{\text{Rif}}$ and the mixed population plated on all three plate types. Populations were allowed to grow and interact for five days before initiating Stage 2, in which populations were manually harvested, and a fixed amount of the liquid resuspension was plated onto agar-casitone plates containing rifampicin, thereby allowing only the $\text{Anc}^{\text{Rif}}$ subpopulations to grow and swarm outward. After four days, $\text{Anc}^{\text{Rif}}$ population sizes at the end of Stage 1 were estimated from Stage 2 swarm sizes. Finally, the negative $\log_{10}$ of the ratio of $\text{Anc}^{\text{Rif}}$ population sizes after competition with Anc clones vs Ev was calculated as a proxy for the relative competitiveness of Ev vs Anc using the reported equation. **c**) Plot showing the relationship between initial population size of inocula of the $\text{Anc}^{\text{Rif}}$ subclones and swarm area after four days of growth and swarming on CTT 1.5% agar with rifampicin. Dots represent the average values of three temporally independent biological replicates ($n = 3$, error bars indicate SEM). **d)** Swarm area relative to the ancestor for all 12 individual evolved populations in their home environments from the same dataset as used for Fig. 2b. The dashed line at 0 represents the ancestral level of swarming on each substrate. The error bars show the SEM for three biological replicates. e) Competitiveness of individual populations relative to the ancestor (dashed line) in their home environment from the same dataset as was used for Fig. 2c. The error bars represent the SEM for three biological assay replicates. **f)** Growth curves estimated from liquid cultures for each population averaged across three replicates and contrasted to the ancestor (dashed black lines). Populations are categorized by evolutionary treatment.

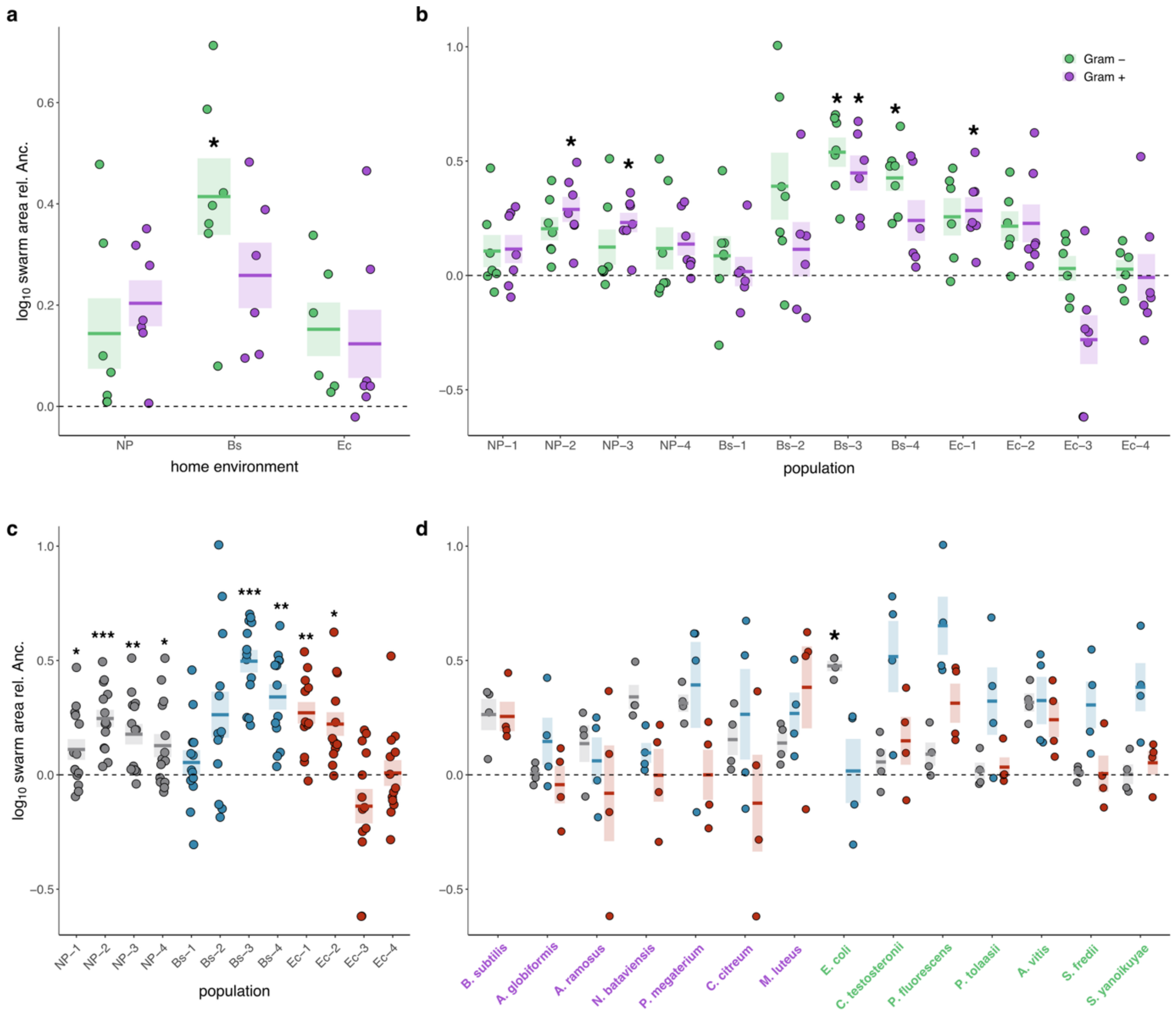


**Supp. Figure 2. Coincidental evolution of swarming rates on foreign prey categorized by prey and Gram type. a**) Average change in swarming rate of evolved treatment population sets (NP, Bs and Ec) on all foreign prey species examined relative to the ancestor (Gram– and Gram+ foreign prey designated by green and purple, respectively; also for panel b). Zero (dashed black line) indicates no evolutionary change (also for panels b, c and d). Each dot represents the treatment-level average swarm size on one prey species. The associated error bars (SE, shaded boxes) reflect the degree to which proportional changes in evolved swarming (relative to the ancestor) vary across prey either Gram– or Gram+ prey species. **b**) The same values shown in panel a for treatment averages are shown for each evolved population. **c)** Average change in swarm size for each population on all foreign prey regardless of Gram-type. Each dot represents the population-level average on one prey species. **d)** Treatment swarm size averages on each prey type relative to the ancestor (dashed line). Each dot represents the evolved population average across three biological assay replicates. For all panels, the shaded area shows the standard error of the mean. Asterisks indicate significant differences between a given average and the ancestral value, all statistical tests are described in the Data Supp. Fig. 2.
*$p < 0.05$ **$p < 0.01$ ***$p < 0.001$

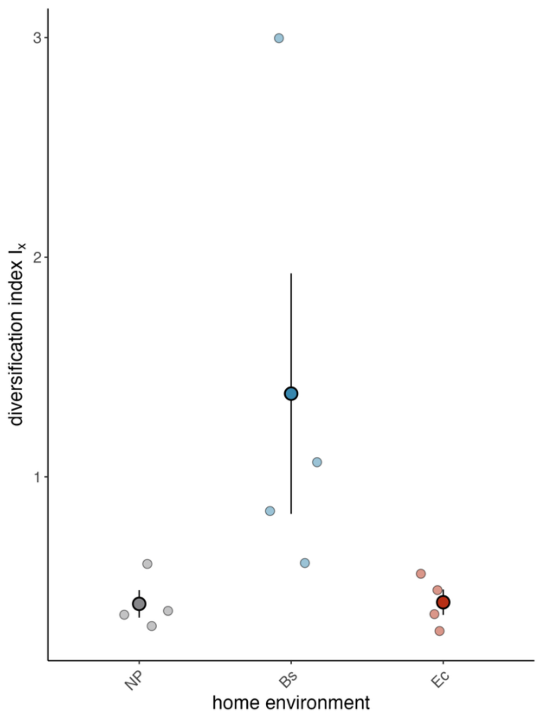


**Supp. Figure 3. Diversification of replicate populations on a panel of foreign prey species.** Diversification index $I_x$ calculated from the variance of each population across 13 (Bs and Ec populations) or 14 (NP populations) foreign prey types adjusted by the average relative swarm area on each prey (following the calculation in Rendueles et al. 2017). The large dots with error bars represent the treatment average with the associated SEM. The Bs population with the largest diversification index is Bs-2.

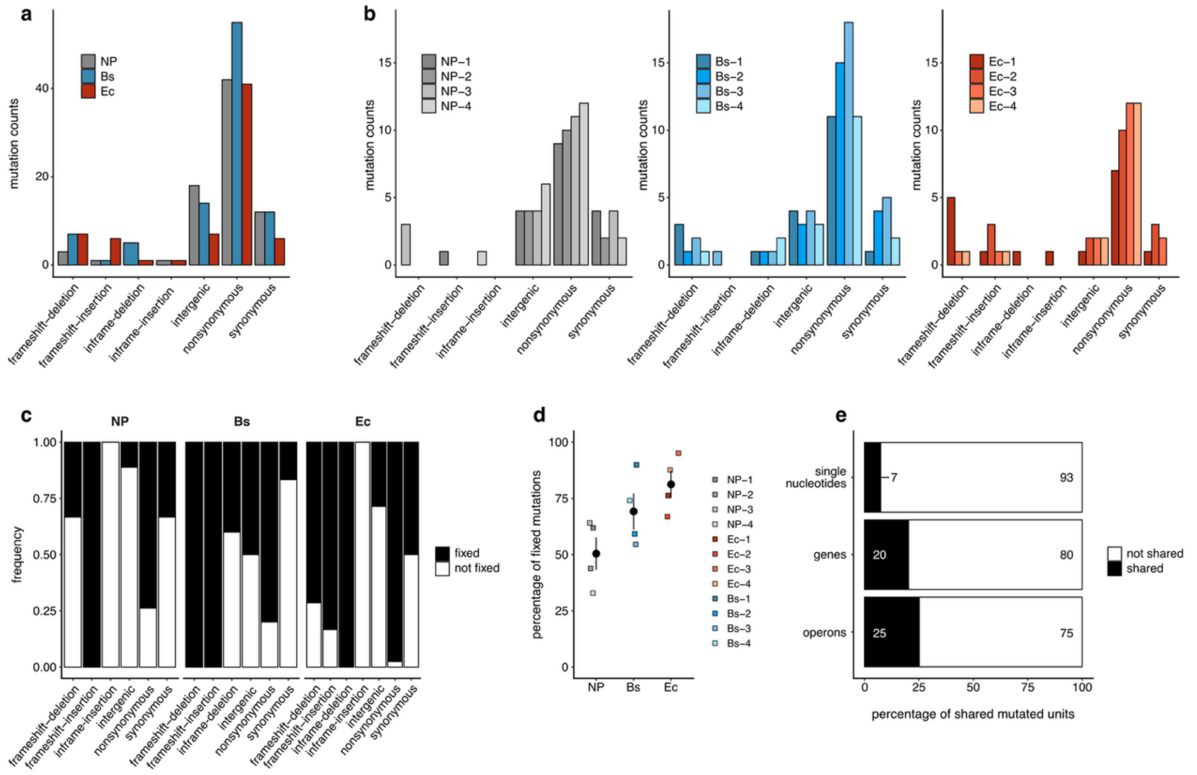


**Supp. Fig 4. Mutation types and type frequencies from whole-population genome sequencing. a**) Total counts of each mutation type found across all population replicates of each evolutionary treatment. Data from the NP, Bs and Ec treatments are shown in grey, blue and red respectively (also for panel b). **b**) Total counts of each mutation type per evolved population across the three evolutionary treatments. **c**) Cross-population frequencies of mutation types for mutations fixed within individual populations for the three evolutionary treatments. **d**) Percentages of all detected mutations that went to fixation within their respective populations, including per-treatment averages. Differently shaded grey, blue, and red squares represent individual population replicates from the NP, Bs, and Ec treatments, respectively. Error bars indicate SEM. **e)** Percentages of individual mutations, mutated genes and mutated operons shared by two or more evolved populations. Numbers within each bar indicate absolute counts of each category.

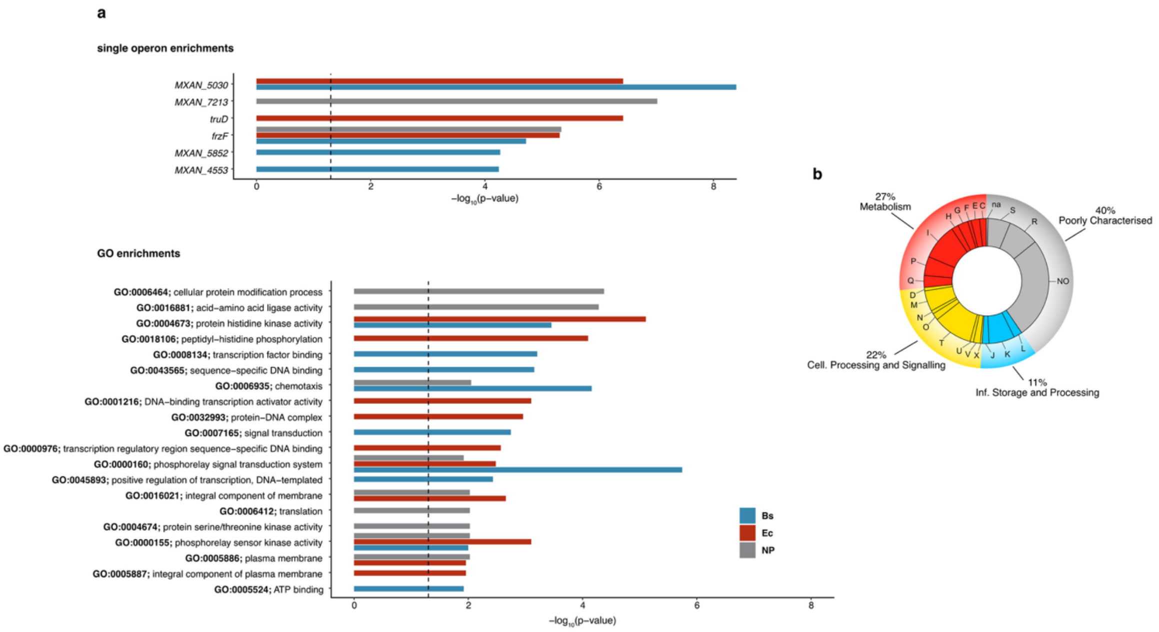


**Supp. Figure 5. Molecular signature of predatory evolution. a**) Top: Operons significantly enriched in mutations in one or more evolutionary treatments (NP in grey, Bs in blue, and Ec in red). Bar length represents the -$\log_{10}$ of the adjusted p-values. The dashed vertical line represents a *p* value of 0.05 (also for GO enrichments below). Bottom: GO categories significantly enriched in mutations in one or more evolutionary treatments. **b**) Percentage distributions of mutated genes across COG classes and subcategories.

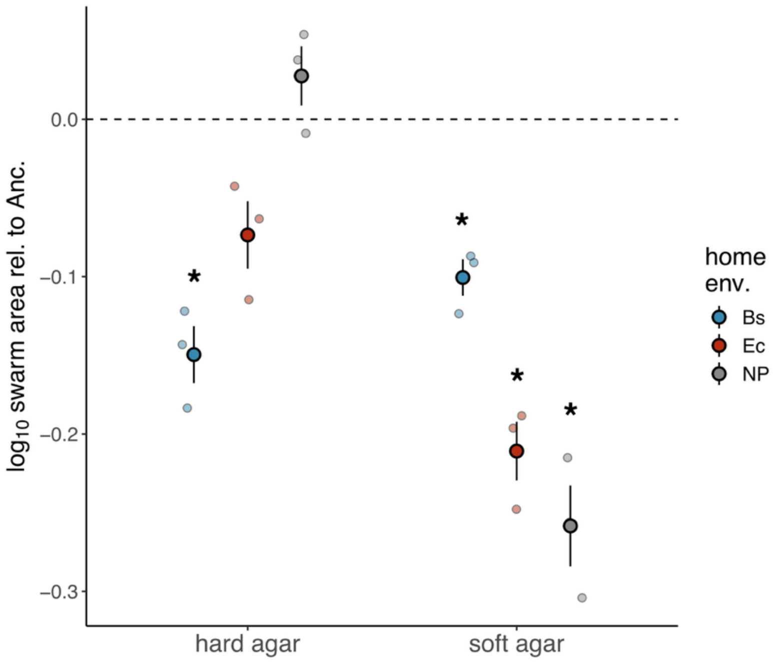


**Supp. Figure 6. Impact of substrate rigidity on swarming.** Treatment-level averages of evolved-population swarming rates on hard and soft CTT agar relative to the ancestor. Averages were calculated from three independent biological replicates ($n = 3$); error bars show the associated SEM. Measurements were taken after four days (instead of seven days as for the predatory swarming measurements), since *M. xanthus* overall swarms faster on soft agar and would reach the edge of the plate before seven days.
* $p < 0.05$

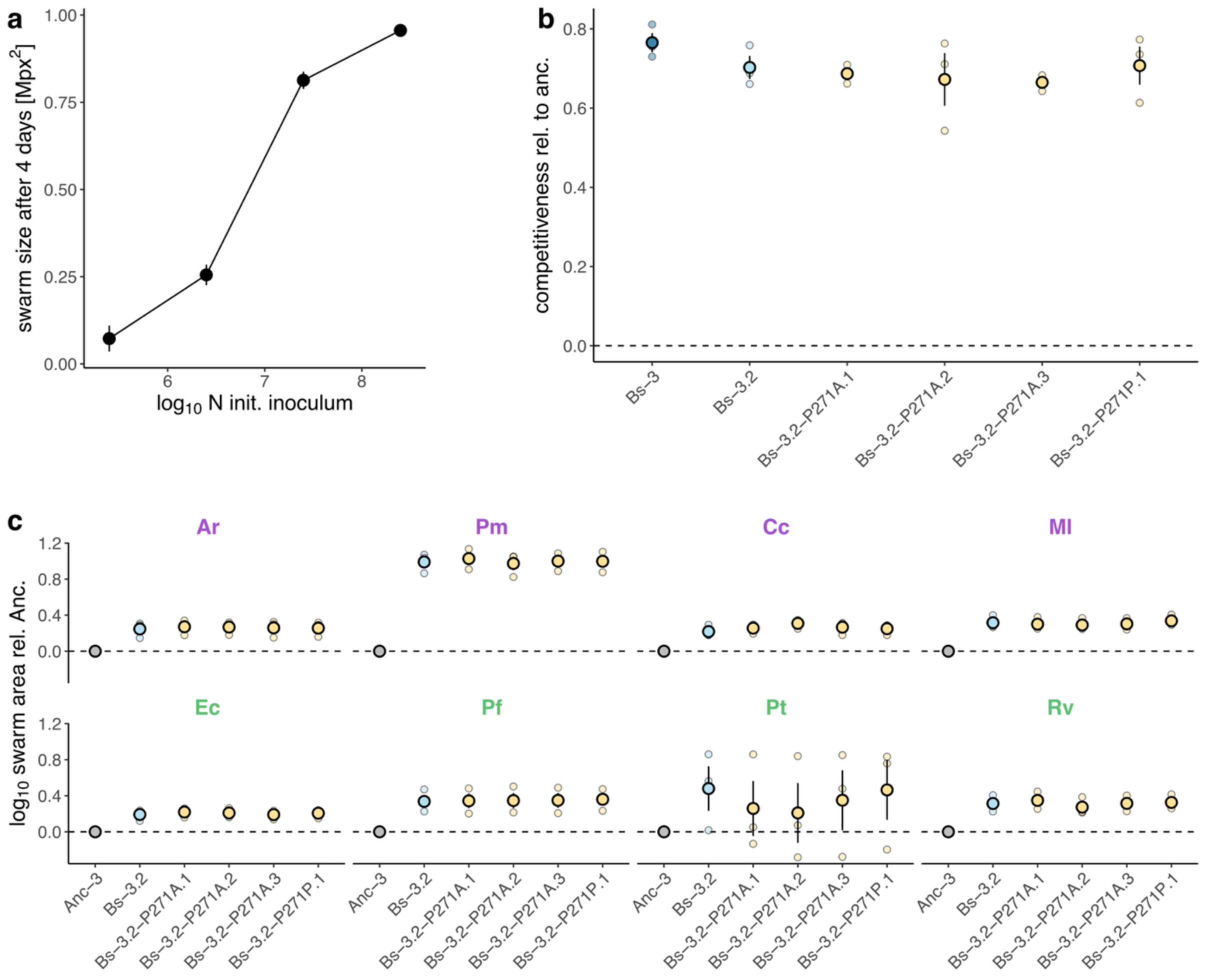


**Supp. Figure 7. Reverting the evolved allele of *MXAN_4699* in clone Bs-3.2 back to the wildtype does not affect competition with the ancestor or swarming on lawns of a panel of prey. a)** Competition of Bs-3 clones from the allele swap with the rifampicin-resistant ancestor of the evolved population on *B. subtilis*. The clones 4699-P271A.1, 4699-P271A.2 and 4699-P271A.3 all have the Bs-3.2 genetic background except are reverted to the ancestral allele of *MXAN_4699*. Clone 4699-P271P underwent the same process for the allele swap but retained the evolved mutation in *MXAN_4699* and serves as a control. The dashed line represents the competition of the rifampicin-resistant and rifampicin-sensitive ancestral clones. **b)** Swarm size after 4 days of the rifampicin-resistant ancestor for colonies started from different inoculation densities. With the relationship between swarm size and starting inoculation density known, swarm size can be used to estimate the population size of the rifampicin-resistant ancestor after competition with another strain. **c)** Predator swarming areas relative to the ancestor of the same strains as in panel (a) (except for population Bs-3) on a set of eight prey other than *B. subtilis*. The top and bottom rows show results on Gram-positive and Gram-negative prey, respectively. The dashed line shows the ancestral level of predatory swarming. As for *B. subtilis*, the presence or absence of the evolved *MXAN_4699* mutation in the Bs-3.2 background did not have an effect on the predator area. Error bars show the SEM for three biological replicates.

**Supp. Table 1: Plasmids, primers and strains used for genetic manipulations.**

| Plasmid | Description | Reference |
|---|---|---|
| pCR-Blunt | Blunt-ended linearised backbone for cloning, kanamycin-resistance | Invitrogen |
| pBJ113 | Plasmid for allele exchange with kanamycin-resistance and *galK* genes for selection & counter-selection | Rodriguez & Spormann 1999 |
| p4699 | pCR-Blunt vector with a 720bp fragment of *MXAN_4699* for knock-in mutants; insertion in *M. xanthus* strains disrupts expression of *MXAN_4699* amplified with GV919 and GV921 | This study |
| p4698 | pCR-Blunt vector with a 1356bp fragment of *MXAN_4698* for knock-in mutants; insertion in *M. xanthus* strains disrupts expression of *MXAN_4698* amplified with GV1147 and GV1149 | This study |
| pCR-4699 | pCR-Blunt vector with a 1029bp fragment of *MXAN_4699* from the ancestral strain (wildtype *MXAN_4699*) amplified with GV1147 and GV1148 | This study |
| pBJ-4699 | pBJ vector with a 1029bp fragment of *MXAN_4699* from the ancestral strain (wildtype *MXAN_4699*) | This study |
| **Primer** | **sequence** | |
| GV1147 | 5'-gatgctcaacaccatgaatcttgaggac-3' | |
| GV1148 | 5'-tcgatggggtcgaagtattcgtact-3' | |
| GV1149 | 5'-aagcttgtcgatgagggacaggtagtacgg-3' | |
| GV919 | 5'-tatcctcaccgccgggagttcatc-3' | |
| GV921 | 5'-ctggaggaaatgctgcgggat-3' | |
| **Strain** | **Description** | **Reference** |
| Bs-3.2 | Clone isolated from the evolved population Bs-3 | This study |
| Bs-3.2::p4698 | Clone with plasmid p4698 insertion, disrupting *MXAN_4698* | This study |
| Bs-3.2::p4699 | Clone with plasmid p4699 insertion, disrupting *MXAN_4699* and *MXAN_4698* | This study |
| Bs-3.2-P271A.1, Bs3.2-P271A.2, Bs-3.2-P271A.3 | Clones after successful allele swap to the wildtype allele, verified by Sanger sequencing | This study |
| Bs-3.2-P271P | Control clone after allele swap which retained the original mutation, verified by Sanger sequencing | This study |

**Supp. Table 2: Mutation types and their frequency in evolved populations.**

| home env. | Bs | | | | Ec | | | | NP | | | | Total |
|---|---|---|---|---|---|---|---|---|---|---|---|---|---|
| population | Bs-1 | Bs-2 | Bs-3 | Bs-4 | Ec-1 | Ec-2 | Ec-3 | Ec-4 | NP-1 | NP-2 | NP-3 | NP-4 | |
| mutations (total) | 20 | 24 | 31 | 19 | 17 | 18 | 18 | 16 | 18 | 16 | 22 | 21 | **240** |
| mutations (polymorphic) | 2 | 10 | 14 | 5 | 4 | 6 | 1 | 2 | 7 | 9 | 8 | 14 | **82** |
| mutations (fixed) | 18 | 14 | 17 | 14 | 13 | 12 | 17 | 14 | 11 | 7 | 14 | 7 | **158** |
| InDels (total) | 4 | 2 | 4 | 3 | 8 | 3 | 2 | 2 | 1 | 0 | 3 | 1 | **33** |
| Intergenic mutations (total) | 4 | 3 | 4 | 3 | 1 | 2 | 2 | 2 | 4 | 4 | 4 | 6 | **39** |
| non-syn. mutations (total) | 11 | 15 | 18 | 11 | 7 | 10 | 12 | 12 | 9 | 10 | 11 | 12 | **138** |
| syn. mutations (total) | 1 | 4 | 5 | 2 | 1 | 3 | 2 | 0 | 4 | 2 | 4 | 2 | **30** |
| non-synonymous (fixed) | 11 | 12 | 10 | 11 | 7 | 9 | 12 | 12 | 8 | 7 | 10 | 6 | **115** |
| Synonymous (fixed) | 1 | 0 | 1 | 0 | 0 | 1 | 2 | 0 | 2 | 0 | 2 | 0 | **9** |
| Target size N (non-synonymous sites at risk) | 5,797,491 | 5,797,421 | 5,797,478 | 5,797,477 | 5,797,495 | 5,797,587 | 5,797,476 | 5,797,467 | 5,797,470 | 5,797,480 | 5,797,474 | 5,797,478 | **mean = 5,797,483** |
| Target size S (synonymous sites at risk) | 2,132,510 | 2,132,489 | 2,132,507 | 2,132,509 | 2,132,514 | 2,132,540 | 2,132,507 | 2,132,503 | 2,132,505 | 2,132,507 | 2,132,507 | 2,132,504 | **mean = 2,132,509** |
| dN/dS (fixed) | 4.0 | >4.4* | 3.7 | >4.0* | >2.6* | 3.3 | 2.2 | >4.4* | 1.5 | >2.6* | 1.8 | >2.2* | **mean= 3.1** |

* For calculations of dN/dS for populations with zero fixed synonymous mutations, we replace zero with an artificial value of 1, generating a minimum-value underestimate of dN/dS, indicated by the '>' sign.

**Supplemental Table 3. MyxoEE-3 cycle-40 evolved populations used in this publication and the corresponding labels used here and for the same populations in Rendueles *et al.* 2015 and Rendueles & Velicer 2017 (see also Rendueles & Velicer 2020 and La Fortezza *et al* 2022).**

| Labels used in this publication | | | Labels used in Rendueles *et al.* 2015 | | |
|---|---|---|---|---|---|
| **Population** | **Ancestor** | **Treatment label & description** | **Population** | **Ancestor** | **Treatment label** |
| NP-1 | Anc-1 | No prey (NP), CTT hard agar (1% casitone) | P1 | GJV1.1 | CTT hard agar (HA) |
| NP-2 | Anc-2 | | P3 | GJV1.2 | |
| NP-3 | Anc-3 | | P5 | GJV1.3 | |
| NP-4 | Anc-4 | | P7 | GJV1.4 | |
| Bs-1 | Anc-1 | *B. subtilis* (Bs), CTT hard agar (1% casitone) with a grown lawn of *B. subtilis* | P97 | GJV1.1 | CTT HA *B. subtilis* |
| Bs-2 | Anc-2 | | P99 | GJV1.2 | |
| Bs-3 | Anc-3 | | P101 | GJV1.3 | |
| Bs-4 | Anc-4 | | P103 | GJV1.4 | |
| Ec-1 | Anc-1 | *E. coli* (Ec), CTT hard agar (1% casitone) with a grown lawn of *E. coli* | P89 | GJV1.1 | CTT HA *E. coli* |
| Ec-2 | Anc-2 | | P91 | GJV1.2 | |
| Ec-3 | Anc-3 | | P93 | GJV1.3 | |
| Ec-4 | Anc-4 | | P95 | GJV1.4 | |
